# Molecular Dynamics simulation of NRAS-Q61 oncogenes and new strategies for *in silico* drug design

**DOI:** 10.1101/2023.05.25.542304

**Authors:** Zheyao Hu, Jordi Marti

## Abstract

The NRAS-mutant subset of melanoma represents the most aggressive and the deadliest types associated with poorer overall survival. Unfortunately, for more than 40 years, no therapeutic agent directly targeting NRAS mutations have been clinically approved yet. Herein, based on microsecond scale molecular dynamics simulations, the concept of NRAS-Q61 mutation classification was firstly proposed. NRAS Q61 positively charged mutations (Q61R and Q61K) was classified together, with a specific targetable pocket, while NRAS-Q61L classified into another category. Moreover, the isomer-sourced structure iteration (ISSI) method was developed for the in silico design of potential inhibitors (HM-516) targeting NRAS mutations. Overall, through this article, we hope to provide the academic and clinical community with new perspective and understanding of NRAS oncoproteins, and propose possible solution to this challenge.

## Introduction

The RAS family is the most frequently mutated oncogene in cancers, such as melanoma, lung and pancreatic cancer.^1^ Clinical data implicated that the most frequently mutated RAS isoform vary by tissue and cancer types.^2^ Whereas, NRAS is the most frequently mutated RAS isoform in melanoma, moreover, the mutated codon of NRAS typically occur at codon 61.^1,2^ As one of the most common cutaneous cancers worldwide, melanoma causes a large number of deaths annually.^3^ And the NRAS-mutant subset of melanoma is aggressive and associated with poorer overall survival. Till now, due to the lack of biological understanding and no strategies of directly targeting NRAS, few specific therapy provided for the NRAS-mutant patients.^4–7^ Therefore, the detailed conformation and local structure of NRAS is urgently needed to be revealed and the discovery of potential drug-like compounds is of great value to the development of directly targeting NRAS mutant therapies.

Drug discovery is an extremely expensive, lengthy and interdisciplinary journey. Usually, an approved drug currently costs several billion and takes more than decade to develop.^8^ In silico drug design can be involved in all stages of drug development, especially in the discovery of high-quality initial lead compounds, which greatly promotes the process and reduces the costs of drug development. Nowadays, with the development of computational resources and methods,^9–13^ and the proliferation of experimental structural data, the trend of using computational physics and molecular modeling for computer-aided drug design has gained enormous momentum, several efficient in silico drug design methods have been released and successfully applied in practice.^14–18^

In this work, benefiting from the advantage that molecular dynamics (MD) simulations can capture the detailed behavior and interaction mechanism of biomolecules in all-atom level with very fine temporal resolution, the impact of NRAS Q61 mutations (wild-type, Q61R, Q61K and Q61L) were firstly investigated. According to the detailed analysis of MD trajectories, NRAS Q61 positively charged mutations (Q61R and Q61K) can be classified into one category with a specific targetable pocket on the surface, while NRAS-Q61L classified into another category. Then we reveal the detailed molecular mechanisms responsible for the polarization of mutated NRAS conformational change behaviors. At the end, combining MD simulations, structural iteration and virtual screening, we propose an isomer-sourced structure iteration (**ISSI**) method for the development of potential lead compounds (**HM-516**) for targeting positively charged NRAS-Q61 mutations. This method fully embodies the idea of structure-based drug design, while cleverly avoids the super-large-scale virtual screening that requires extremely high computing resources.

## Results and discussion

In this section, first, we investigated the conformational changes of the four main isoforms of NRAS-Q61 in aqueous ionic solution. The only difference between these four NRAS is located at codons 61, with the sequences of the four isoforms shown in Fig. 8 of "Supporting Information". Then radial distribution functions (*g_AB_*(*r*)) and potential of mean force (*W_AB_*(*r*)) were employed to reveal the mechanisms corresponding to the conformational changes of the different NRAS-Q61 mutants. Finally, we demonstrate the process of developing potential lead compounds (HM-516) for NRAS mutations using the isomer-sourced structure iteration (ISSI) method. Atomic detail sketches of the main residues described in this part are reported in Fig. 9. All other remaining important data & figures have been reported in SI.

### The fluctuations and stability of NRAS and its mutated isoforms

Firstly, we employed Root Mean Square Deviations (RMSD) and Root Mean Square Fluctuations (RMSF) to investigate the conformational fluctuations and stability of NRAS-WT and its three mutant isoforms (Fig. 1). The RMSD are defined by Eq.(1):

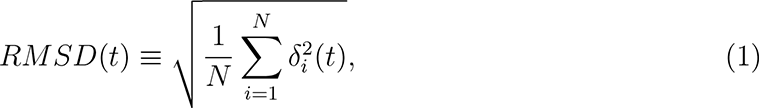

**Figure 1:**
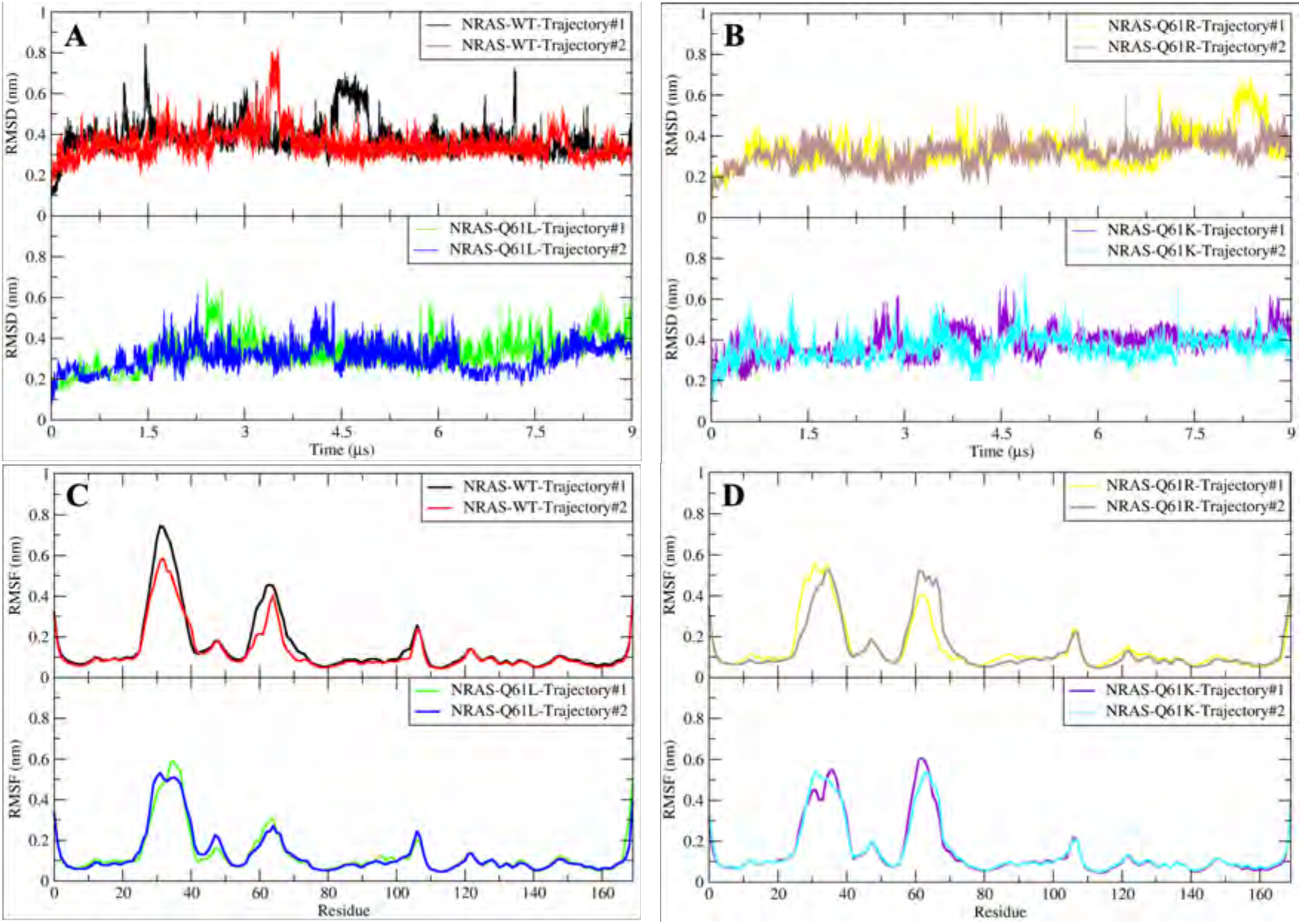
The RMSD and RMSF of different NRAS isoforms. (A) The RMSD of NRAS-WT and NRAS-Q61L; (B) The RMSD of NRAS-Q61R and NRAS-Q61K; (C) The RMSF of NRAS-WT and NRAS-Q61L; (D) The RMSF of NRAS-Q61R and NRAS-Q61K.

where *δ_i_* is the difference in distance between the atom *i* (located at *x_i_*(*t*)) of the catalytic domain and the equivalent location in the crystal structure. Further, RMSF are defined by Eq.(2):

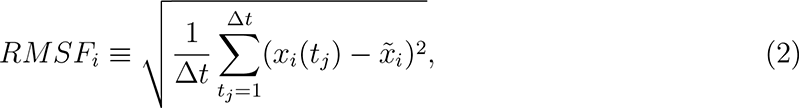

Where 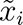 is the time average of *x_i_* and Δ*t* is the time interval where the average has been taken. RMSD results show the fluctuations and stability of the conformations of the proteins. As shown in Fig. 1 (A) NRAS-WT case, the RMSD value fluctuated around 0.36 nm steadily, and there were several transient conformational fluctuations (RMSD ∼0.6 nm) during the whole simulation. Unlike KRAS-WT which has a distinct and persistent conformational fluctuation, such that "State-I" and "State-II" can be detected easily,^19^ NRAS-WT is more likely to maintain conformational fluctuations in a relatively small range. In a similar fashion, whether Q61 mutated to positively charged amino acids (ARG and LYS, case of mutations Q61R and Q61K) or mutated to non-polar amino acid (LEU, case of mutation Q61L), the fluctuation of the RMSD value are similar to those of NRAS-WT. From the perspective of residues, RMSF revealed their flexibility during the full simulation. This feature is similar to the widely studied high conformational flexibility regions Switch-I (SW-I) and Switch-II (SW-II) of KRAS,^19–22^ also detected in NRAS protein case, as shown in Fig. 1 (C-D). In NRAS-WT case the flexibility of the SW-I domain is around 1.5 to 2 fold larger than that of SW-II. Noticeably, when Q61 was mutated to positively charged amino acids, the flexibility of SW-I decreased slightly, whereas the RMSF values of SW-II increased leading to similar flexibilities of SW-I and SW-II. Interestingly, when Q61 was mutated to a non-polar amino acid, the trend of RMSF was opposite to that of positively charged mutations. The flexibility of the SW-II domain decreased around 1-fold compare with the NRAS-WT and NRAS-Q61 positively charged mutations. At the same time, it is worth noting that the peak of NRAS-Q61L SW-II tends to the direction of increasing residue sequence. For example, the RMSF value of NRAS LEU61 decreased significantly, and meanwhile the peak of SW-II shifted to the right side. The above RMSD and RMSF information suggests that: (1) NRAS proteins are more structurally stable than KRAS proteins over the timescales involved in this paper; (2) There exist different mechanisms in conformational changes of different NRAS isoforms; (3) It corroborates that to a large extent, the NRAS proteins are similar to other RAS family proteins, and their conformational changes are mainly embodied by SW-I and SW-II.

### The effect of Q61 mutations on the dominant conformations of NRAS

According to the evidences provided by RMSD and RMSF analysis and in order to investigate the differences in the conformational changes of NRAS-WT and its mutants in more detail, we displayed and analysed the Gibbs free energy profiles of NRAS proteins. Gibbs free energy profiles are of high significance to characterize the dominant conformations of target proteins during simulation. ^23^ In this case, such analytical tools will allow us to directly track the effects of GLN61 mutations on the conformational changes of the NRAS. Since the main movement of the protein is concentrated in SW-I, SW-II and the residues with RMSF value greater than 0.2 nm we chose to compute Gibbs free energy landscapes by using two specific variables such as RMSD and radius of gyration (*R_g_*), as defined in Eq.3:

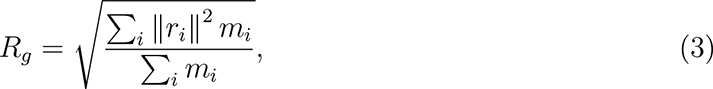

where *m_i_* is the mass of atom *i* and *r_i_* the position of the same atom with respect to the centre of mass of the selected group. The method employed to obtain the free energy profiles has been the so-called "Principal Component Analysis",^24,25^ where the components RMSD and *R_g_* worked as reaction coordinates (Figure 2 A1, B1, C1 and D1). To corroborate the full convergence of the free energy landscapes reported in Fig.2, we report in SI (Fig. 10 to Fig. 13) the contribution of the two independent MD trajectories that we have employed in this work (A,B), compared with their average (C), as shown in Figure 2 A1, B1, C1 and D1.

**Figure 2:**
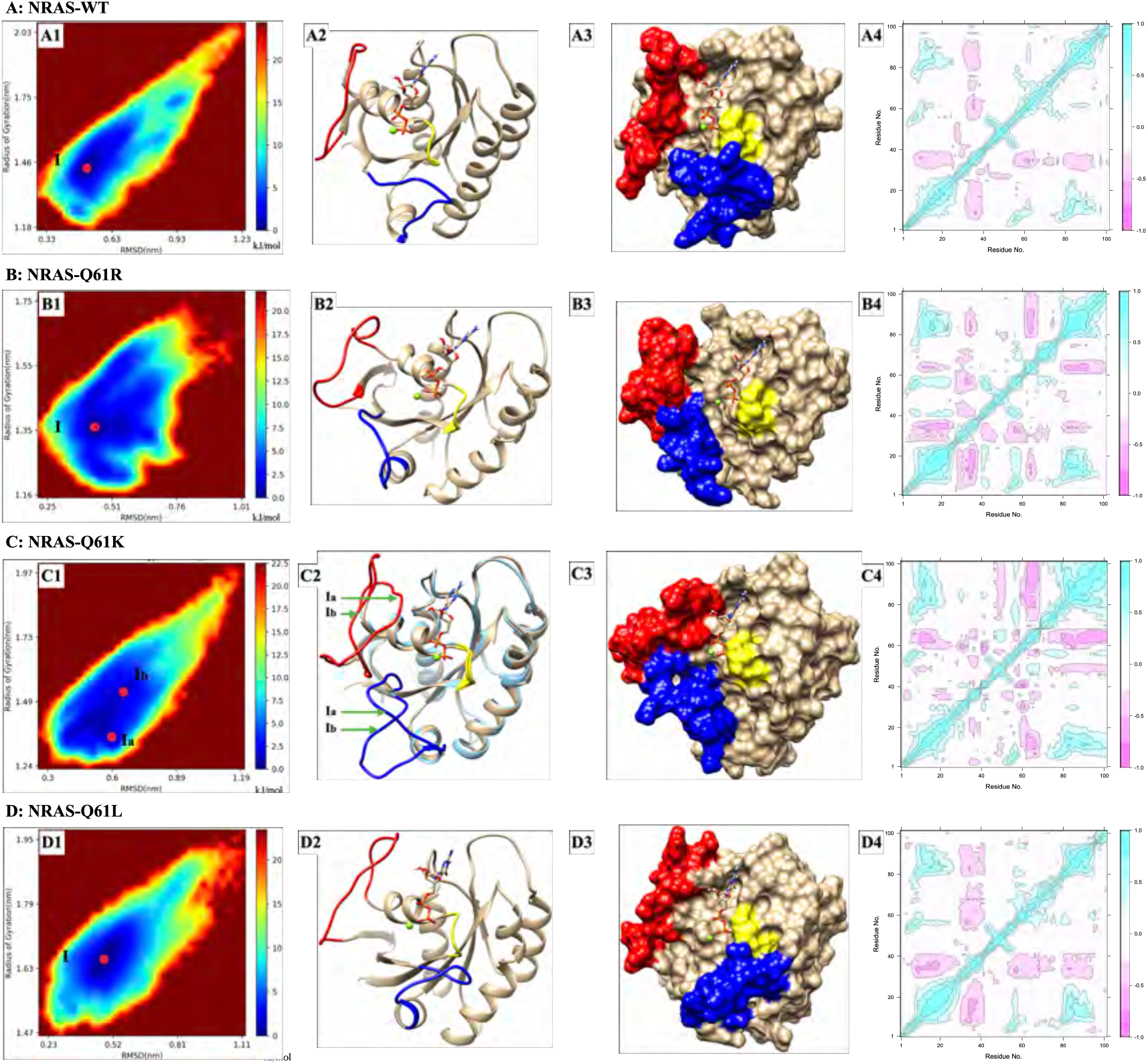
Trajectory analysis of NRAS isoforms and their representative dominant conformations. (A1) The "Gibbs Free-Eenergy Landscape" of NRAS-WT; (A2) The representative dominant conformation of NRAS-WT basin I; (A3) The surface of representative dominant conformation of NRAS-WT basin I; (A4) The “Residue Cross Correlation” analysis of NRAS-WT. (B1) The "Gibbs Free-Eenergy Landscape" of NRAS-Q61R; (B2) The representative dominant conformation of NRAS-Q61R basin I; (B3) The surface of representative dominant conformation of NRAS-Q61R basin I; (B4) The “Residue Cross Correlation” analysis of NRAS-Q61R. (C1) The "Gibbs Free-Eenergy Landscape" of NRAS-Q61K; (C2) The representative dominant conformations of NRAS-Q61K basins Ia and Ib; (C3) The surface of representative dominant conformation of NRAS-Q61K basin Ia; (C4) The “Residue Cross Correlation” analysis of NRAS-Q61K. (D1) The "Gibbs Free-Eenergy Landscape" of NRAS-Q61L; (D2) The representative dominant conformation of NRAS-Q61L basin I; (D3) The surface of representative dominant conformation of NRAS-Q61L basin I; (D4) The “Residue Cross Correlation” analysis of NRAS-Q61L. Red: SW-I; Blue: SW-II; Yellow: P-loop.

In the NRAS-WT case, there mainly exists one free energy basin (I), Figure 2 A1. The Gibbs free energy basin (I) represent the main stable state of NRAS-WT during MD simulations. Like the dominant conformation (Figure 2 A2 and A3) obtained from the MD trajectories according to the free energy basin I, SW-I is slightly open and SW-II tends to be close to the P-loop and *α*-helix 3. From this point of view, the conformational changes of NRAS-WT are similar to the fluctuations in the state I of KRAS-WT.**^?^** In order to better characterize the trajectory variation of NRAS-WT, we further analyzed the MD trajectory of NRAS-WT using the “Cross-Correlation Analysis” tool in "Bio3D",^26^ see Figure 2 A4 "Residue cross correlation" of NRAS-WT. "Residue cross correlation" can help us to better understand the trend of protein’s residue-to-residue distance changes during MD simulations, we depicted in pink color those amino acid residues that are far away each other and in turquoise blue color those residues that are close to each other. As we can see form Figure 2 A4, the SW-I exhibited the trend of moving away from the SW-II and the main structure of the NRAS-WT protein, while SW-II behaved differently: SW-II showed a tendency to be closer to P-loop and *α*-helix 3, a fact that agrees well with the results shown on Figure 2 A2 and A3. The mutations of NRAS-Q61 to positively charged side chains (Q61R and Q61K), showed some significant differences with NRAS-WT. As for NRAS-Q61R, one single Gibbs free energy basin was also detected, Figure 2 B1. In this case, the representative conformation of basin I (Figure 2 B2 and B3) is quite different from the representative dominant conformation of NRAS-WT. We can clearly monitor that the behavior of SW-II in NRAS-Q61R structure is opposite to that of NRAS-WT, the SW-II tends to stay away from the main structure of the NRAS-Q61R (such as the P-loop and *α*-helix 3). In this case, SW-II and SW-I tend to be close to each other and their surfaces in close contact. Interestingly, after the separation of SW-II from the main structure of NRAS-Q61R, the generated cavity on the surface of NRAS-Q61R may serve as a targetable pocket for drug design, see Figure 2 B2 and B3. Finally, the trend of protein’s residue-to-residue distance changing during the simulation was also verified by the "Residue cross correlation" analysis of NRAS-Q61R, Figure 2 B4. From Figure 2 B4, we can clearly see that the tail of SW-I (residues ∼ 39 to 40) and the head amino acid of SW-II (residues ∼ 58 to 60) tended to be close to each other, whereas the tail of SW-II (residues ∼ 61 to 70) showed a clear tendency to move away beyond the main body of NRAS (P-loop and *α*-helix 3). In NRAS-Q61K case, in a similar fashion as in mutation Q61R, there exist a single and extended free energy basin, which is divided into several small energy basins, with Ia and Ib as examples (See Figure 2 C1). Basin Ia is the one with lowest free energy (set to 0 kJ/mol) and it has been chosen as reference so that basin Ib shows a barrier of 0.4 kJ/mol, when transitions from basin Ia to Ib are considered. These basins are separated by such low free energy barriers that interconversion between the two sub-basins is well within the energy of thermal fluctuations. The differences between basins Ia and Ib are mainly on SW-I and SW-II, with the SW-I conformation of Ib being slightly counterclockwise compared to Ia and the SW-II conformation of Ib being more open than that of Ia (Figure 2 C2). The surface of the representative dominant conformation Ia of NRAS-Q61K exhibited a similar style compared with NRAS-Q61R, so that it displays a potentialy targetable pocket for pharmacological purposes (see Figure 2 C3 and B3). The "Residue cross correlation" results of NRAS-Q61K (Figure 2 C4) showed a similar style compared to NRAS-Q61R. The tendency of the tail of SW-I and the head of SW-II to be in close contact is weakened compared with NRAS-Q61R, whereas the tail of SW-II showed a clear tendency to move apart from the main body of NRAS (P-loop and *α*-helix 3). When Q61 of NRAS was mutated to the non-polar amino acid LEU (Q61L), the Gibbs free energy landscape showed a marked single basin (see D1 in Figure 2). As it can be seen from the representative crystal structure and its surface version of this particular free energy basin I (Figure 2 D2 and D3), this dominant conformation of NRAS-Q61L corresponds to SW-I slightly open, with SW-II separated separated from SW-I and tightly combined with the P-loop and *α*-helix 3 regions.

To further characterize the effect of mutation of Q61 on NRAS protein, we provided the “Trajectory Distribution Scatter Plot” comparison of NRAS and its Q61 mutations, see Figure 3. Here we also chose to compute “Trajectory Distribution Scatter Plot” by using two specific variables RMSD and radius of gyration (*R_g_*), and each point in it represents a frame in the MD trajectories. Figure 3A shows the trajectory distribution of NRAS-WT, as the reference for comparison to its Q61 mutations. Combined with the Gibbs free energy analysis in Figure 2, it can be seen that the coordinates of the most densely distributed NRAS-WT trajectories were (0.5, 1.45). When the Q61 mutated to LEU, the coordinates of the most densely distributed trajectories is (0.5, 1.65), and the overall trajectories showed a tendency to distribute towards the direction of *R_g_* increasing (see Figure 3B). In the NRAS-Q61R case, the coordinates of the most densely distributed trajectories were (0.45, 1.35), and the overall trajectories showed a tendency to reduction of both *R_g_* and RMSD, when compared to WT (see Figure 3C). Finally, for the NRAS-Q61K case, the overall trajectory coincided with NRAS-WT, but the coordinates of the most densely distributed trajectory had a tendency to move towards the direction of decreasing *R_g_* and increasing RMSD (see Figure 3D). According to the present analysis we found that the Q61 mutations had effects on the conformational distribution of NRAS proteins. When Q61 is mutated to a positively charged amino acid, the conformational distribution area of the protein is roughly unchanged, but the *R_g_* parameters of dominant conformation tend to decrease; on the contrary, when Q61 is mutated to a non-polar amino acid, the *R_g_* of the protein will increase. This helped us better classify the different mutations of NRAS.

**Figure 3:**
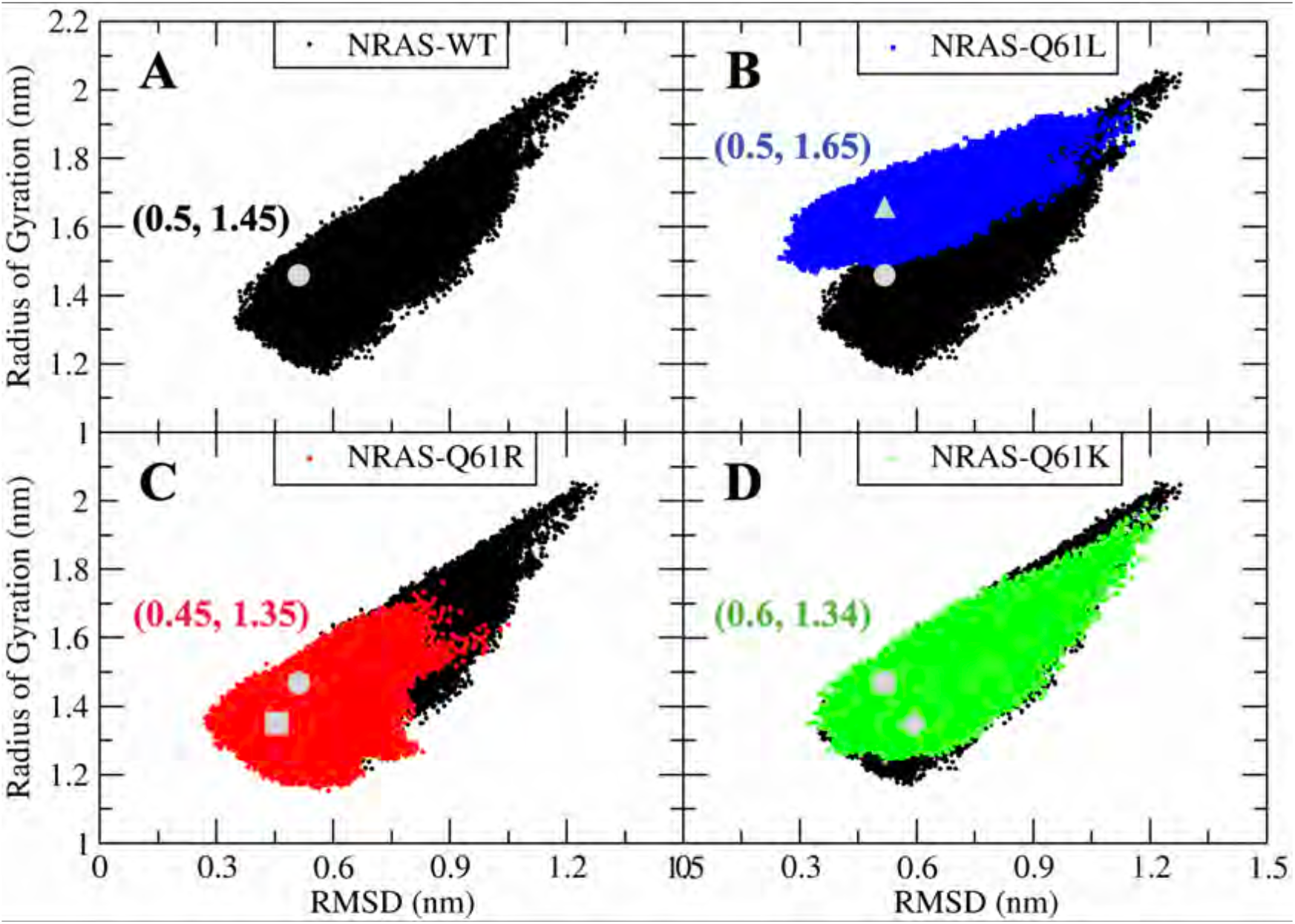
The “Trajectory Distribution Scatter Plot” comparison of NRAS and its Q61 mutations. (A) NRAS-WT as the reference (black), the dominant conformational coordinates are (0.5, 1.45); (B) NRAS-WT with NRAS-Q61L (blue), the dominant conformational coordinates of NRAS-Q61L are (0.5, 1.65); (C) NRAS-WT with NRAS-Q61R (red), the dominant conformational coordinates of NRAS-Q61R are (0.45, 1.35); (D) NRAS-WT with NRAS-Q61K (green), the dominant conformational coordinates of NRAS-Q61K are (0.6, 1.34).

### The different mechanisms driving conformational changes of NRAS

The above study shows that different types of mutations in Q61 position can have different effects on the conformational changes of NRAS. ARG and LYS are positively charged side chain amino acids such that they can be grouped into the same class. We have seen that after mutation of Q61, NRAS-Q61R and NRAS-Q61K show similar conformational changes, SW-I and SW-II tend to be close each other and at the same time SW-II moves away from the P-loop and *α*-helix 3 and creates a potentially targetable pocket between SW-I/SW-II and the P-loop/*α*-helix 3. Another group is the non-polar side chain amino acid LEU61 mutation. A comparison of the latter to the wild-type and positively charged mutation groups showed that the fluctuation of SW-II was significantly reduced and that SW-II could be tightly combined with P-loop and *α*-helix 3. Here, we provide two molecular mechanisms that could lead to the generation of the two distinct mutant groups described above. First, we considered the so-called atomic pair radial distribution functions *g_AB_*(*r*) (Eq.(4)) to explore the changes in the ability to form hydrogen bonds (HB) with water molecules after the mutation of Q61, as reported in Figure 4.

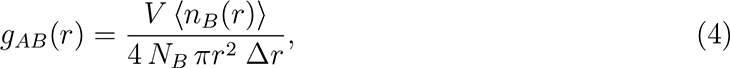

**Figure 4:**
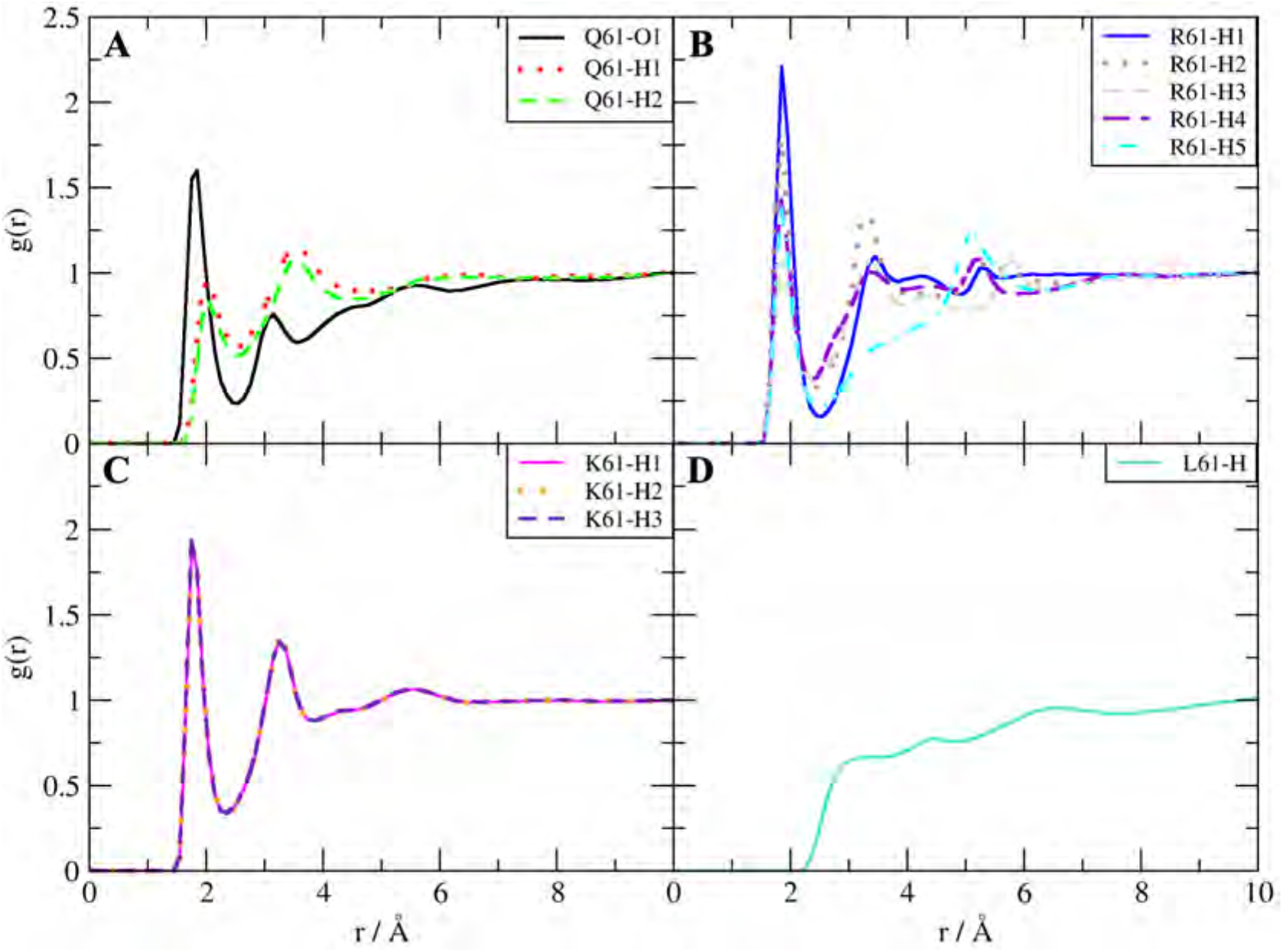
Radial distribution function analysis of the 61st amino acid of NRAS and water. (A) NRAS-WT case; (B) NRAS-Q61R case; (C) NRAS-Q61K case; (D) NRAS-Q61L case.

where *n_B_*(*r*) is the number of atoms of species *B* surrounding a given atom of species *A* inside a spherical shell of width Δ*r*. *V* is the total volume of the system and *N_B_* is the total number of particles of species *B*. As a general feature, the active sites of wild type NRAS Q61 side chain capable of forming HB are oxygen "O1", hydrogens "H1" and "H2" (see Fig. 9). O1 is able to bind hydrogens of water molecules and the HB length is around

1.85 Å, typical of bonds formed by biomolecules and water^27^ (see Figure 4A). The ability of hydrogens "H1" and "H2" to form HB with the solution is weaker than that of "O1", given the lower height of the RDF (Figure 4A), with a typical HB length of 2.05 Å. In NRAS-Q61R case, the active sites of R61 side chain capable of forming HB are five hydrogen atoms "H1" to "H5", see Fig. 9. As shown in Figure 4B, these active hydrogens can interact with oxygens of water molecules with a characteristical HB lenght of 1.85Å, with the first maximum of the RDF enhanced in comparison with the NRAS-WT case. This means that when Q61 is mutated to R61, the interaction of R61 with aqueous solution is enhanced. Similar to NRAS-Q61R, when Q61 was mutated to LYS, we observed three active hydrogens forming HBs with solvating water (see Fig. 9) and the HB interaction of K61 with water molecules was also enhanced, with the HB length of 1.75 Åand the first maximum of the RDF around *g*(*r*) = 1.92 (See Figure 4C). Interestingly, the RDF of the three hydrogens almost coincide. Contrary to wild type and proteins Q61R and Q61K (both with positive net charge), the mutation Q61L (with a non polar LEU site) showed no HB at all (see Figure 4D), indicating strong hydrophobic characteristics.

In order to further quantitatively analyze the degree of enhanced HB interaction with aqueous solution when Q61 mutated to positively charged amino acid, we consider the radial distances between two species as our order parameters to calculate the potential of mean force (PMF) through the so-called reversible work *W_AB_*(*r*), as defined by Eq.(5):

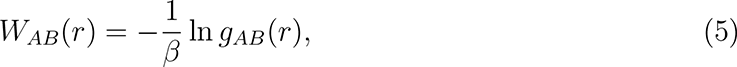

where *β* = 1*/*(*k_B_T*) is the Boltzmann factor, *k_B_*the Boltzmann constant and T the temperature. We show PMF for wild type, Q61R and Q61K NRAS proteins in Fig. 15. A free-energy barrier defined by a neat first minimum and a first maximum of *W* (*r*) is clearly seen in all three cases. As expected, the locations directly match the first maxima of the corresponding RDF. And the full set of positions and free-energy barriers for the three cases has been reported in Table 1. There, we can observe overall barrier of NRAS-WT is 3 *k_B_T*, which correspond to 1.8 kcal/mol (1 *k_B_T* = 0.616 kcal/mol). For the NRAS-Q61R and NRAS-Q61K their free-energy barriers are 8.6 *k_B_T* (5.3 kcal/mol) and 5.4 *k_B_T* (3.3 kcal/mol), respectively. From the above results, we can clearly see that while Q61 mutated to the positively charged amino acids (such as ARG and LYS), the HB interaction of this site with aqueous solution is enhanced by ∼ 2 to 3 fold, which may be one of the mechanisms driving the dissociation of SW-II from protein surfaces.

**Table 1:** Free-energy barriers *δW* (in *k_B_T*) from reversible work calculations for the interaction between 61st amino acid side chains and water. In order to quantify the height of all barriers, 1 *k_B_T* = 0.616 kcal/mol. *δW* here is the average *δW* after the average of Trajectory#1 and Trajectory#2.

| Mutants | Site 1 | Site 2 | $r_{min}(\text{Å})$ | $r_{max}(\text{Å})$ | $\Delta W$ (in $k_B T$) |
| --- | --- | --- | --- | --- | --- |
| 3*Wild Type | O1 | H (water) | 1.85 | 2.55 | 1.9 |
|  | H1 | O (water) | 2.05 | 2.55 | 0.6 |
|  | H2 | O (water) | 2.05 | 2.55 | 0.5 |
| 5*Q61R | H1 | O (water) | 1.85 | 2.55 | 2.7 |
|  | H2 | O (water) | 1.85 | 2.45 | 1.7 |
|  | H3 | O (water) | 1.85 | 2.45 | 1.0 |
|  | H4 | O (water) | 1.85 | 2.35 | 1.3 |
|  | H5 | O (water) | 1.85 | 2.55 | 1.9 |
| 3*Q61K | H1 | O (water) | 1.75 | 2.35 | 1.8 |
|  | H2 | O (water) | 1.75 | 2.35 | 1.8 |
|  | H3 | O (water) | 1.75 | 2.35 | 1.8 |

For the non-polar side chain amino acid LEU61 mutation, the mechanism of NRAS-Q61L conformational changes is quite different from the positively charged mutations. A hydrophobic pocket on the surface of the *α*-helix 3 was detected, see Figure 5. This hydrophobic pocket is mainly composed of the hydrophobic structures (see labeled green parts in Fig. 9) of ASP92, LEU95, TYR96 and GLN99 side chains. During the dynamic evolution of NRAS-Q61L, this hydrophobic pocket can capture the side chain of L61 and, simultaneously, the hydrogen atom "H1" on the amino group of the main chain of L61 can form a stable HB interaction with the "O1" of TYR96, which further stabilizes L61 and promotes the tight binding of SW-II to the protein surface. This mechanism well explains the difference in conformational changes between NRAS-Q61L and other NRAS isoforms. In order to better characterize this special hydrophobic pocket of L61, we also displayed the time-dependent atomic group–group distances between selected amino acid residue side chains and the time-dependent atomic site–site distances between LEU61(H1) and TYP96(O1), see Fig. 16 and Fig. 17.

**Figure 5:**
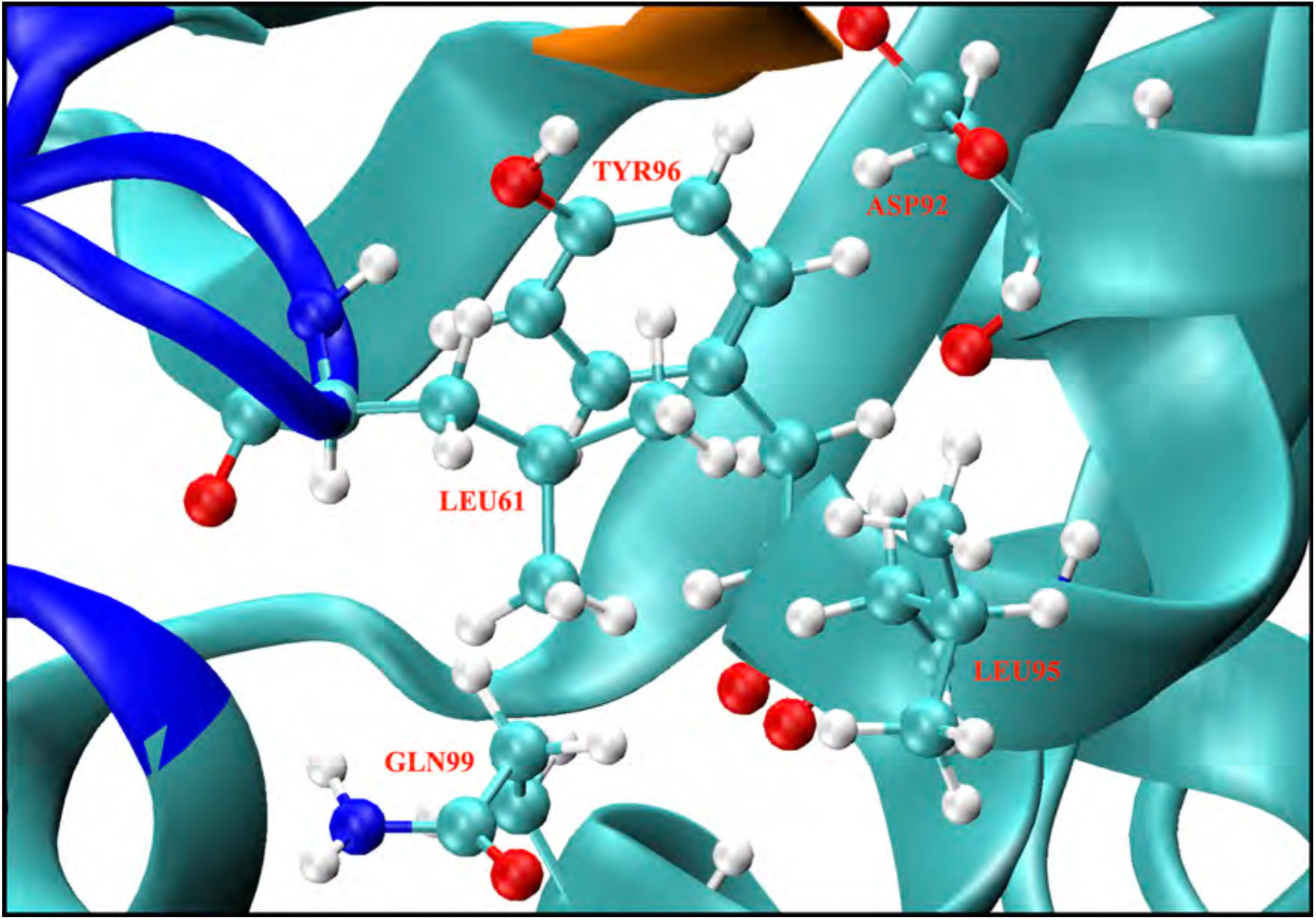
The detected hydrophobic pocket for LEU61.

### *In silico* development of potential inhibitors targeting NRAS Q61 mutations using the ISSI method

In the previous section, we classified the differences in conformation changes after mutation of NRAS Q61 in detail. When the Q61 mutated to positively charged amino acids (R and K), there exists a significant channel on the surface of NRAS. Here, for the sake of convenience, we take the targetable pocket on the surface of NRAS-Q61R as an example to carry out corresponding in silico drug design, structure evolution and screening, hoping to provide potential inhibitor template for targeting mutated NRAS proteins. Informed by the MRTX1133,^28^ see Fig. 19, the highly efficient noncovalent KRAS-G12D inhibitor targeting the Switch-II pocket (similar to the targetable pockets on the mutated NRAS), we designed a series of compounds with a pyrido[4,3-*d*]pyrimidine scaffold in silico with the anticipation that after the structure evolution and screening could found the potential candidate compounds could effectively bind to the surface pocket of mutated NRAS. During the isomer-sourced structure iteration (**ISSI**) process, we mainly carried out 3 rounds of structural iterations, and finally obtained the potential inhibitor **C6** (**HM-516)**) of NRAS-Q61R (detailed structural iteration process, see supporting information).

Further, considering the: ligand/receptor flexibility, charge distribution, and surrounding water & ions molecules, well designed MD simulations (4 independent trajectories, 10 **HM-516** + NRAS-Q61R, total 10 *µ*s) were employed to explore in detail the dynamic binding mechanism of **HM-516** to the NRAS-Q61R surface pocket. The results of MD simulations revealed in detail the active site of **HM-516** and the interaction mode between **HM-516** and NRAS-Q61R, and also verified the predicted interaction mode during ISSI structure iteration.

Here we introduce the main binding mechanism (Fig. 6) and multiple interaction modes and dynamic movies are included in more detail in the supporting material. The guanine module of **HM-516** acts like the "bridge", limiting the mobility of SW-I and fixing it around the *α*-helix 1 through through hydrogen bonding with SER17, ASP30 and coordination with the GDP-Mg^2+^ complex. Similar to the guanine module, the DBD part of **HM-516** plays a button-like role, mainly forming hydrogen bond interactions with SW-II of NRAS. The pyrrolizidine ring located on C2-position of the pyrido[4,3-*d*]pyrimidine core acts like boat anchor and can anchor the hydrophobic pocket located inside NRAS through hydrophobic interactions. At the same time, the hydrogen bonds interaction between ARG68 and O/N atoms (Fig. 6, shown by the cyan right-angle dotted line) further fix the pyrrolizidine ring. Finally, the naphthyl amine substituents further enhances the affinity of **HM-516** to NRAS through the *π*-*π* stacking with TYR64 and the hydrogen bond interaction between the amino group and LEU95 & GLN99. Overall, the combination of the three generations of structural iteration, docking analysis and MD simulation led to the discovery of **HM-516**, a potential selective inhibitors of NRAS Q61 positively charged mutations.

**Figure 6:**
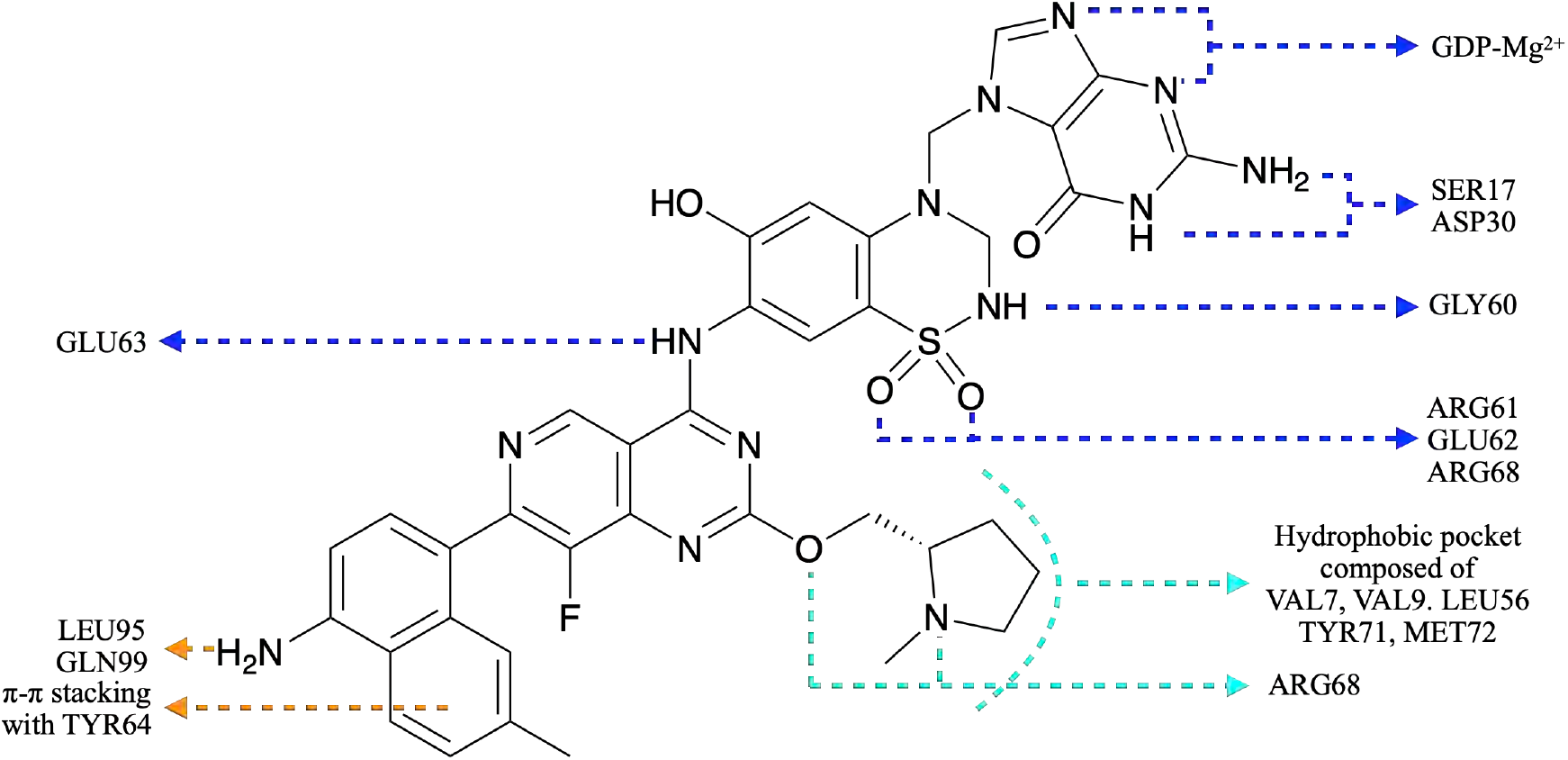
Dynamic binding model of HM-516 to the targetable pocket of NRAS-Q61R

## Conclusions

NRAS mutations typically occur at codon 61, there mainly exists the following 4 mutations: Q61R, Q61K, Q61H and Q61L. Since the proportion of NRAS-Q61H mutation in all NRAS-Q61 mutations is very low, so the wild-type, Q61R, Q61K and Q61L were investigated in detail in this paper. Through extensive MD simulation and trajectory analysis, the conformational change rules of NRAS proteins and the underlying molecular mechanisms were revealed. Mutations at codon Q61 result in altered charging at this position, which in turn alters the interaction of SW-II with its surroundings. When the Q61 was mutated to the positively charged amino acid (e.g. R and K), the HB interaction of the amino acid residue at position 61 with aqueous solution will be enhanced. Compared with the wild type, the HB interactions are enhanced by 3.3 kcal/mol and 1.4 kcal/mol, respectively. In a sense, this enables the aqueous solution to dissolve SW-II better, and thereby stripping SW-II off the protein surface. In contrast, when Q61 is mutated to the non-polar amino acid LEU, the affinity of aqueous solution to SW-II decreases, and the side chain of LEU can be captured by the hydrophobic pocket on the *α*-helix 3, thereby anchoring SW-II in the protein surface. These two opposing molecular mechanisms lead to two distinct classifications of NRAS mutations. NRAS Q61 positively charged mutations (Q61R and Q61K) were classified into one category with a specific targetable pocket on their surface, while NRAS-Q61L classified into another category. Given that the proportion of positively charged mutations in NRAS Q61 is greater than 80%, then an isomer-sourced structure iteration (**ISSI**) method was proposed to design the potential inhibitors targeting the targetable pocket of the NRAS positively charged mutation group. In section 3.4 of the article, taking NRAS-Q61R as an example, the whole process of designing and obtaining the potential inhibitor **HM-516** through the **ISSI** method is fully demonstrated. Furthermore, combined with MD simulation, the interaction mode between **HM-516** and NRAS-Q61R was explored and disclosed at all-atom level. In conclusion, we have disclosed the conformational changes of NRAS oncoproteins in detail at the atomic level and proposed a new concept of NRAS mutation classification for the first time. Moreover, in silico drug design method "isomer-sourced structure iteration" was proposed for the discovery of potential inhibitors of NRAS oncoproteins.

## Methods

### Molecular dynamics simulation parameter settings

Our main computational tool has been microsecond scale molecular dynamics (MD). In MD, after the choice of reliable force fields the corresponding Newton’s equations of motion are integrated numerically,^29^ allowing us to monitor each individual atom in the system in a wide varierty of setups, including liquids at interfaces in solid walls or biological membranes among others.^30,31^ We fixed the number of particles, the pressure and the temperature of the system, while the volume fluctuated accordingly to the barostat employed. MD can model hydrogens at the classical^32,33^ or quantum levels^34^ and, in addition to energetic and structural properties, and it provides information on free-energy landscapes and access to time-dependent quantities such as the diffusion coefficients or spectral densities,^35^ enhancing its applicability. In the present work we conducted MD simulations of four NRAS isoforms with sequences represented in Fig. 8. Each system contained one isoform of the NRAS-GDP complex fully solvated by 5,699 TIP3P water molecules**^?^** in potassium chloride solution at the human body concentration(0.15 M) and magnesium chloride solution concentration(0.03 M) yielding a system size of 19,900 atoms. All MD inputs were generated using CHARMM-GUI solution builder^36–38^ and the CHARMM36m force field.^39^ The force field used also includes the parameterization of the species GDP (it can be searched as “GDP” in the corresponding CHARMM36m topology file: https://www.charmm-gui.org/?doc=archive&lib=csml) All bonds involving hydrogens were set to fixed lengths, allowing fluctuations of bond distances and angles for the remaining atoms. Crystal structure of GDP-bound NRAS proteins was downloaded from RCSB PDB Protein Data Bank, ^40^ file name "6zio". The four sets of NRAS proteins (wild type, Q61R, Q61K and Q61L) were solvated in a water box, all systems were energy minimised and well equilibrated (NVT ensemble) before generating the production MD. Eight independent production runs were performed within the NPT ensemble for 9 *µ*s (total 72 *µ*s). All meaningful properties were either averaged from the two runs. The pressure and temperature were set at 1 atm and 310.15 K respectively, in order to simulate the human body environment. In all MD simulations, the GROMACS/2021 package was employed.^41^ Time steps of 2 fs were used in all production simulations and the particle mesh Ewald method with Coulomb radius of 1.2 nm was employed to compute long-ranged electrostatic interactions. The cutoff for Lennard-Jones interactions was set to 1.2 nm. Pressure was controlled by a Parrinello-Rahman piston with damping coefficient of 5 ps*^−^*^1^ whereas temperature was controlled by a Nosé-Hoover thermostat with a damping coefficient of 1 ps*^−^*^1^. Periodic boundary conditions in three directions of space have been taken. We employed the “gmx-sham” tool of the GROMACS/2021 package to performed the Gibbs free energy landscape analysis. Other alternatives such as transition path sampling^42–44^ or metadynamics^45,46^ were not considered here because of their high computational cost for large systems. The MD simulation parameters for subsequent exploration of the binding mode of **HM-516** and NRAS-Q61R are the same as above, except that the number of water molecules is 15535 and the MD inputs were generated using CHARMM-GUI multicomponent assembler, in order to simulate 10 molecules of **HM-516** together with mutated NRAS in water box.

### Trajectory analysis and visualization

The GROMACS/2021 package, ^41^ the software VMD,^47^ UCSF Chimera^48^ and R-package "Bio3D"^26^ were used for trajectory analysis and visualization.

### Isomer-sourced structure iteration (ISSI)

This section provides an overview of current isomer-sourced structure iteration (ISSI), including input content preparation, structure iteration and judging criteria. And the ISSI drug discovery process follows the established procedure shown schematically in Fig. 7. The input content mainly consists of “receptor structures" and “isomer-sourced template”. Here, we use crystal structure (PDB id 6zio) as the initial structure, the MD simulation was employed to obtain the "receptor structure" for the further drug design process. In this method the MRTX-1133^28^ was selected as the isomer-sourced template molecule for subsequent structural iterations, the 3D models of the template molecule and subsequent iteration molecules were constructed by CgenFF.**^? ?^** Then the docking analysis (Fig. 7, Docking-I^49^) between the selected seeds (e.g. MRTX-1133) and receptor explored their binding pattern, providing direction and guidance for subsequent structural iterations. In this method, the number of structural iterations can be increased or decreased as needed. In this article, we set up three generations of structural iterations based on the initial docking analysis (Fig. 7, Docking-I), corresponding to Fig. 20, Fig. 22 and Fig. 24. During each structural iteration, the corresponding compound library will be generated, and the most potential candidate will be screened out according to the screening rules to enter the next generation of structural iteration. Here two judgment criteria are used to measure the potential of candidate compounds: (1) the docking score between the candidate molecule and the receptor, (2) the number of hydrogen bonds formed between the candidate molecule and the receptor. Moreover, the selected molecules will be further verified by MD simulation or experiment.

**Figure 7:**
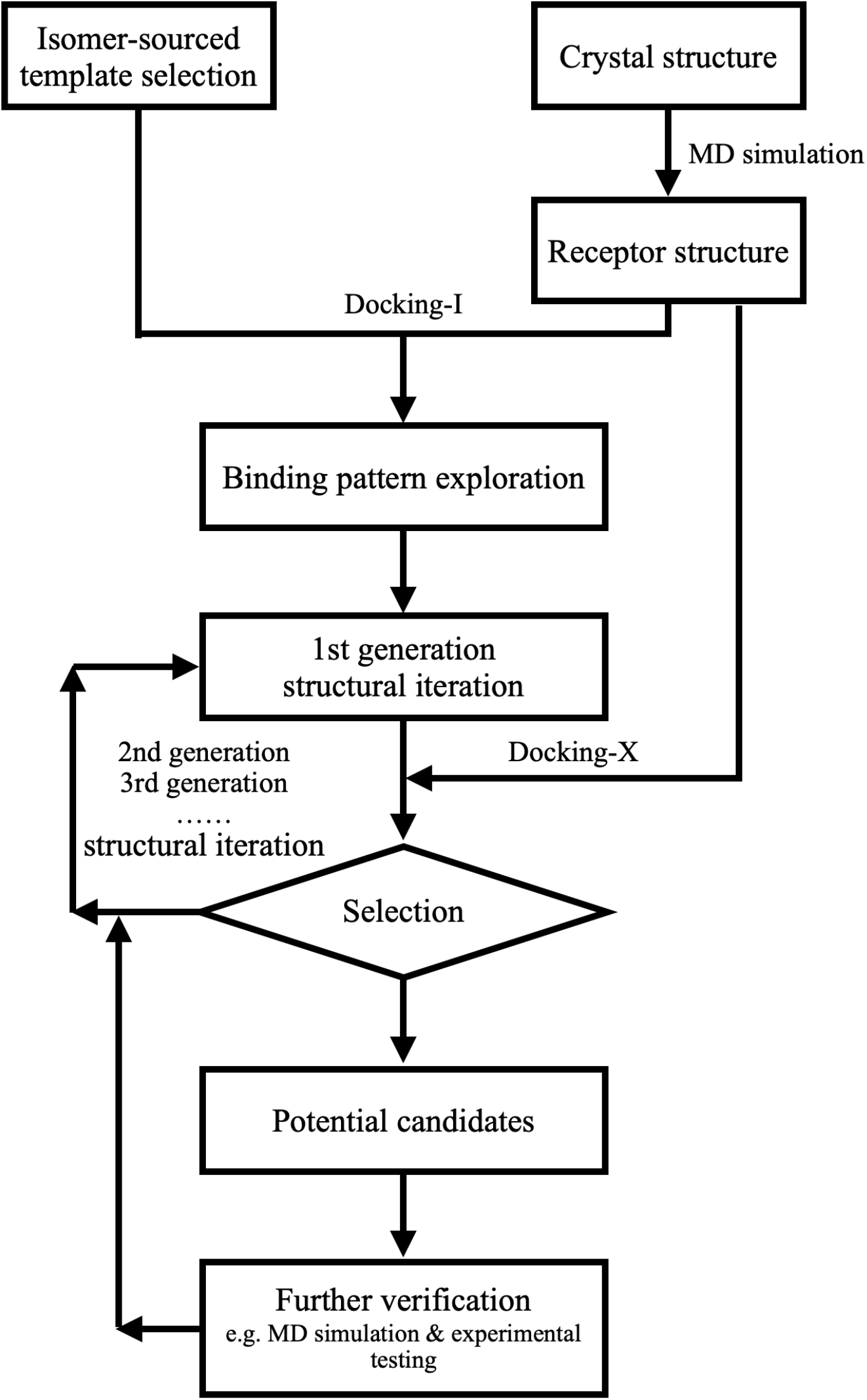
Schematic diagram of the "Isomer-sourced structure iteration" *in silico* drug design method.

## Supporting information

### Initial setups, data and convergence of the simulations

The four classes of NRAS proteins that have been considered in this work are reported in Fig. 8, where the initial structure of NRAS-WT, was acquired from PDB bank (6zio.pdb). In order to make clear the atomic sites that will be described and analysed in the article, we represent sketches of the main amino acid structures analysed in the present work in Fig. 9.

**Figure 8:**
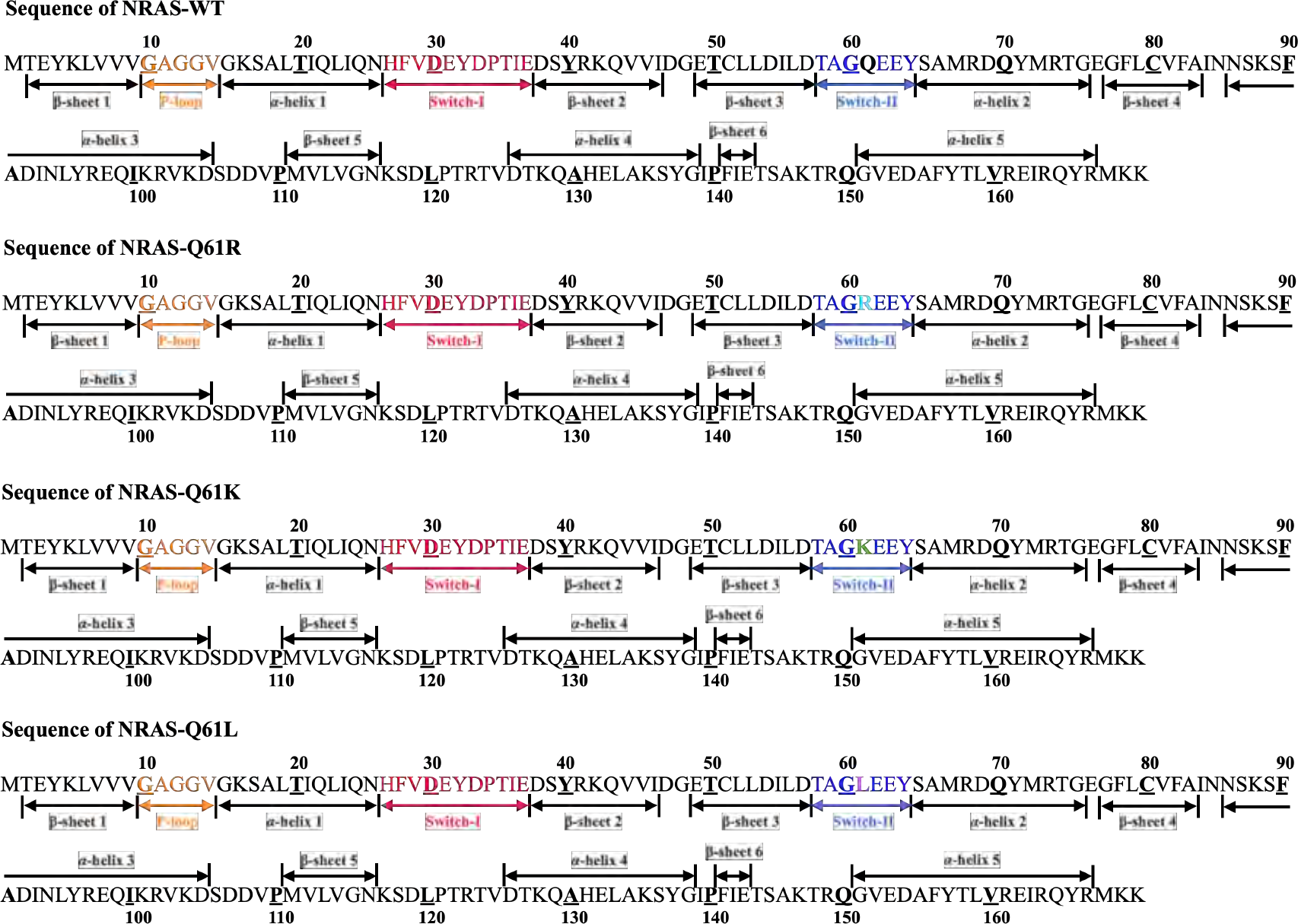
The sequence of four NRAS isoforms: SW-I (red), SW-II (blue), Q61 (black), R61 (cyan), K61 (green) and L61 (magenta).

**Figure 9:**
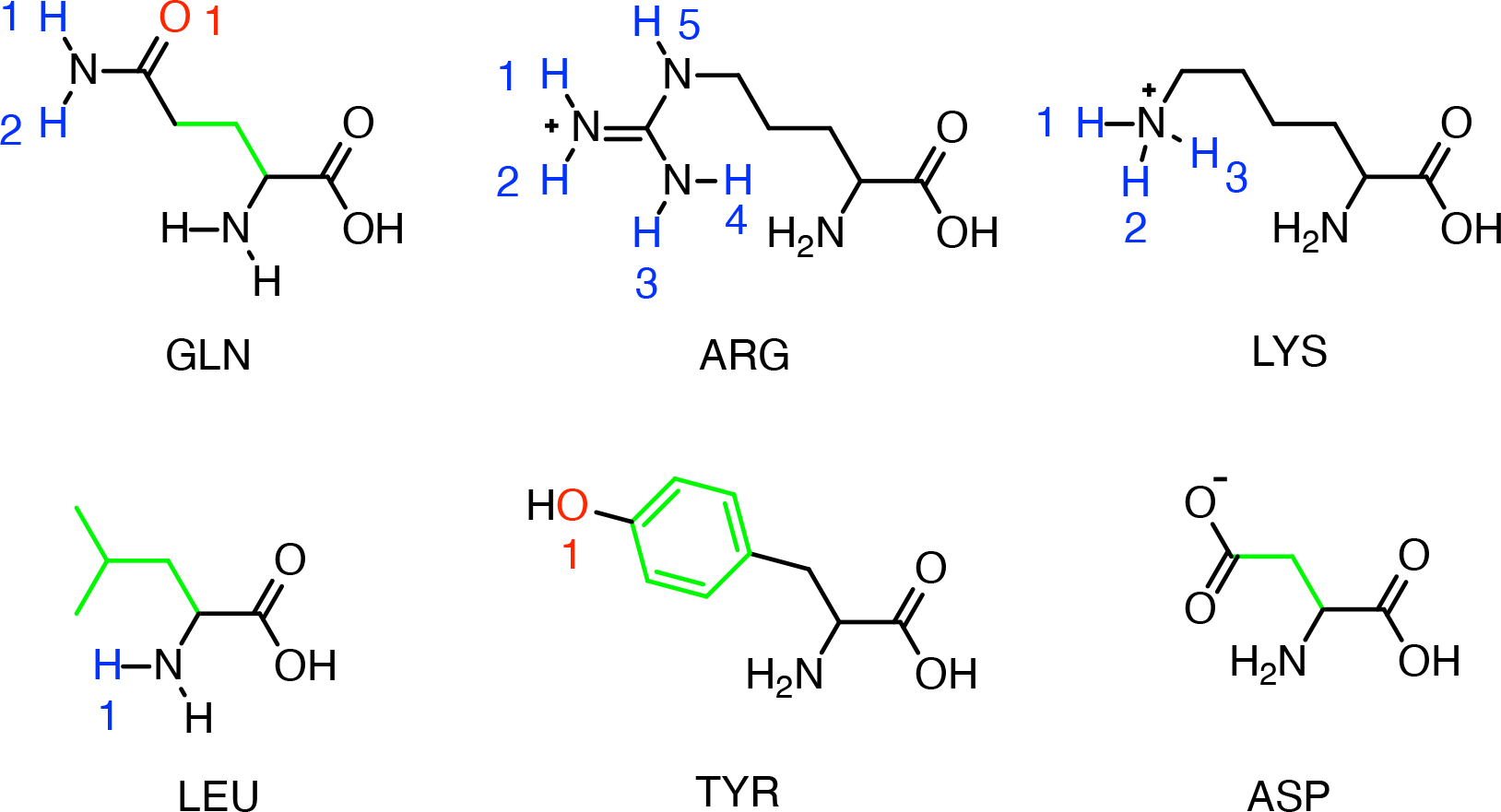
Sketches of amino acid residues of NRAS protein mentioned in the text.

The equilibration of our simulation systems has been reached after a few tens of nanoseconds. Afterwards, eight long independent trajectories of 9 *µ*s each have been produced for exploring the conformational changes of NRAS and its mutants. In order to show the convergence of our results, we are reporting the calculation of the free-energy surface corresponding to the four NRAS isoforms case from the two runs (A, B) and its average (C) as shown from Fig. 10 to Fig. 13. We have included a superposition (D) of the stable state configuration of NRAS-WT and its mutations for the three sets reported (A,B,C).

**Figure 10:**
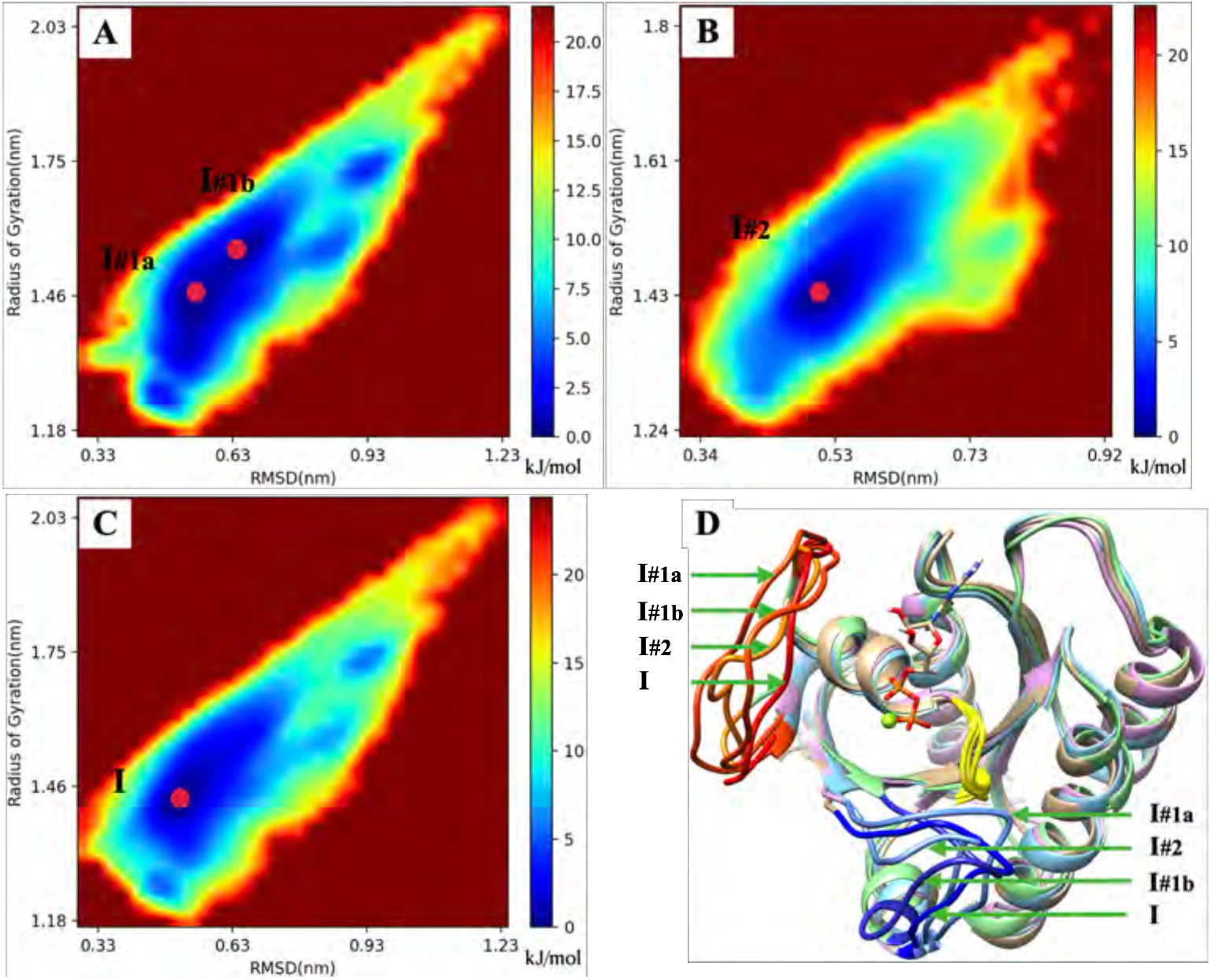
The "Gibbs Free Energy" analysis of NRAS-WT; (A) The “Gibbs Free-Energy Landscape” of NRAS-WT Trajectory#1; (B) The “Gibbs Free-Energy Landscape” of NRAS-WT Trajectory#2; (C) The “Gibbs Free-Energy Landscape” of NRAS-WT (The average of Trajectory#1 and Trajectory#2); (D) The comparison of the most dominant conformations of “Average”, “Trajectory#1” and “Trajectory#2”. Red: Switch-I (Average), Orange Red: Switch-I (Trajectory#1), Orange: Switch-I (Trajectory#2), Blue: Switch-II (Average), Cornflower Blue: Switch-II (Trajectory#1), Medium Blue: Switch-II (Trajectory#2), Yellow: P-loop, Green: GDP-bound Mg^2+^. Water and other ions have been hidden.

**Figure 11:**
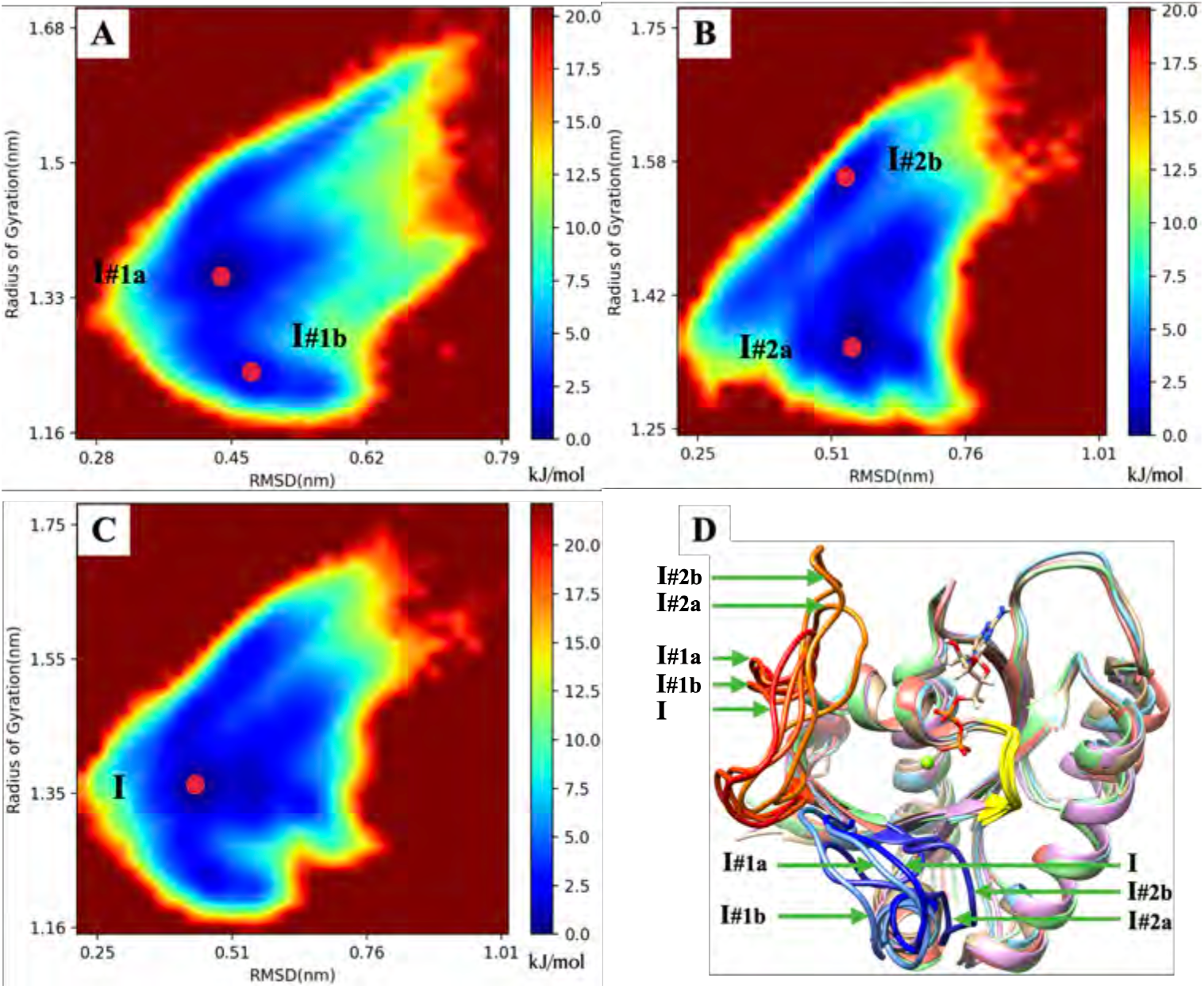
The "Gibbs Free Energy" analysis of NRAS-Q61R; (A) The “Gibbs Free-Energy Landscape” of NRAS-Q61R Trajectory#1; (B) The “Gibbs Free-Energy Landscape” of NRAS-Q61R Trajectory#2; (C) The “Gibbs Free-Energy Landscape” of NRAS-Q61R (The average of Trajectory#1 and Trajectory#2); (D) The comparison of the most dominant conformations of “Average”, “Trajectory#1” and “Trajectory#2”. Red: Switch-I (Average), Orange Red: Switch-I (Trajectory#1), Orange: Switch-I (Trajectory#2), Blue: Switch-II (Average), Cornflower Blue: Switch-II (Trajectory#1), Medium Blue: Switch-II (Trajectory#2), Yellow: P-loop, Green: GDP-bound Mg^2+^. Water and other ions have been hidden.

**Figure 12:**
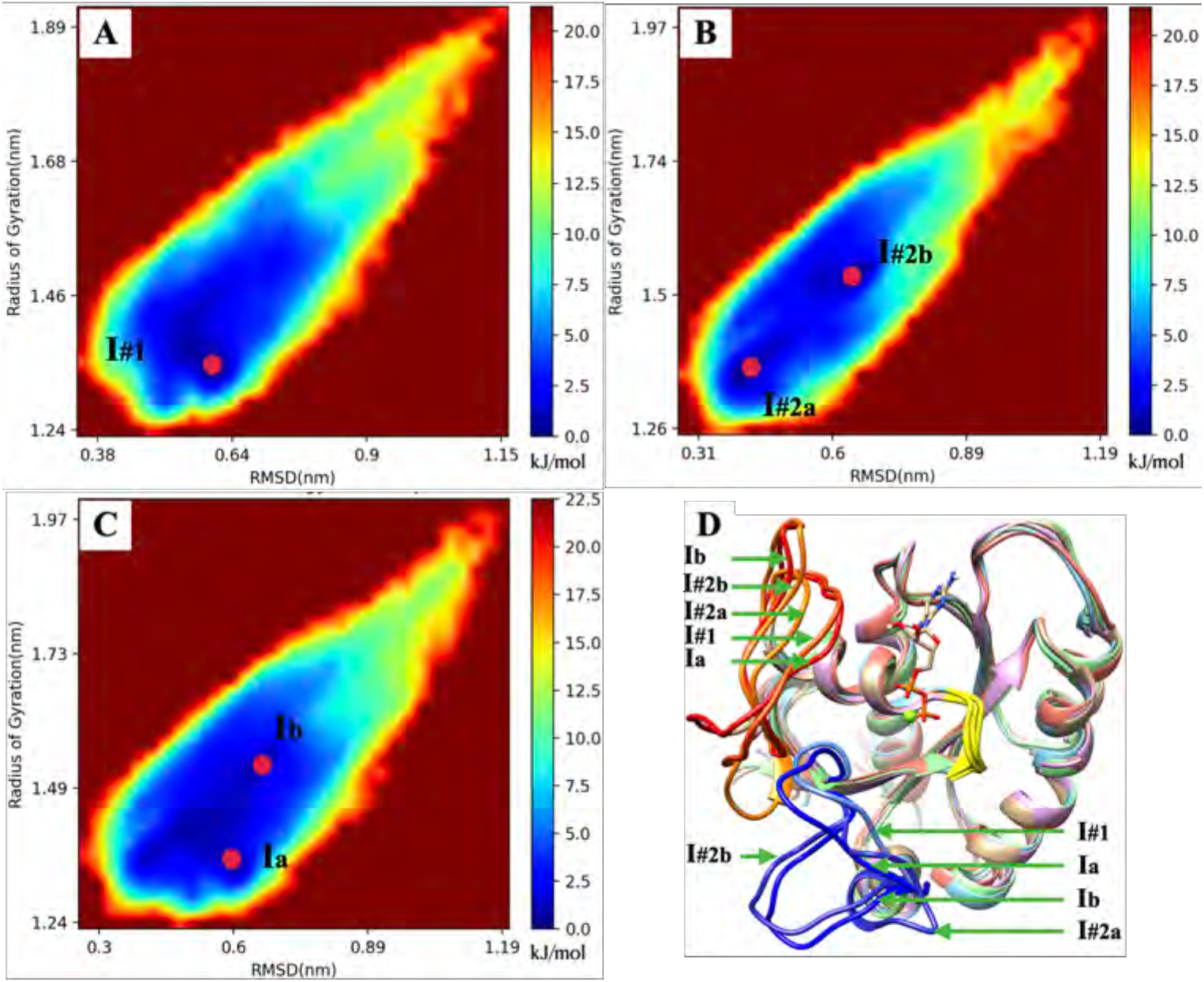
The "Gibbs Free Energy" analysis of NRAS-Q61K; (A) The “Gibbs Free-Energy Landscape” of NRAS-Q61K Trajectory#1; (B) The “Gibbs Free-Energy Landscape” of NRAS-Q61K Trajectory#2; (C) The “Gibbs Free-Energy Landscape” of NRAS-Q61K (The average of Trajectory#1 and Trajectory#2); (D) The comparison of the most dominant conformations of “Average”, “Trajectory#1” and “Trajectory#2”. Red: Switch-I (Average), Orange Red: Switch-I (Trajectory#1), Orange: Switch-I (Trajectory#2), Blue: Switch-II (Average), Cornflower Blue: Switch-II (Trajectory#1), Medium Blue: Switch-II (Trajectory#2), Yellow: P-loop, Green: GDP-bound Mg^2+^. Water and other ions have been hidden.

**Figure 13:**
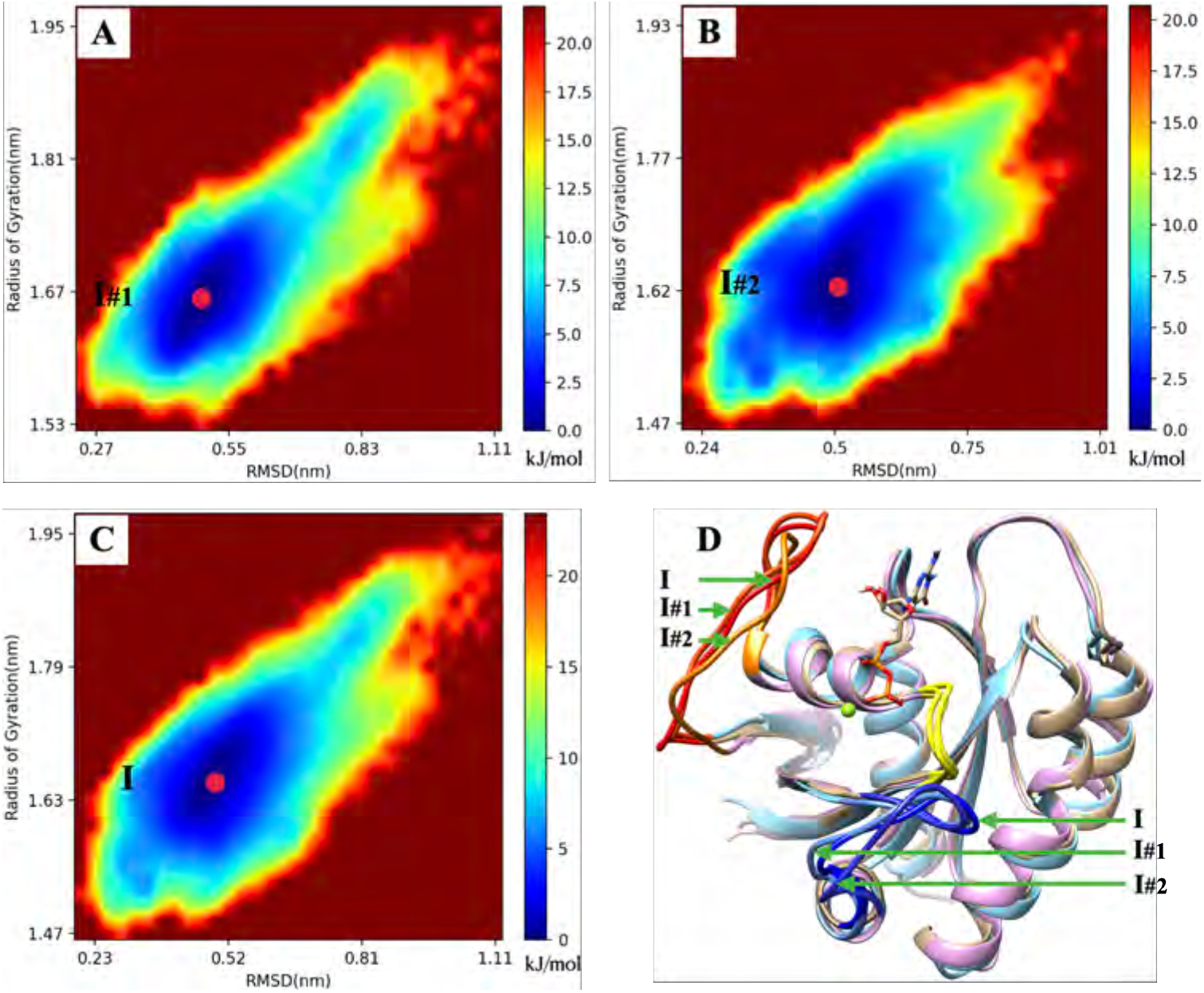
The "Gibbs Free Energy" analysis of NRAS-Q61L; (A) The “Gibbs Free-Energy Landscape” of NRAS-Q61L Trajectory#1; (B) The “Gibbs Free-Energy Landscape” of NRAS-Q61L Trajectory#2; (C) The “Gibbs Free-Energy Landscape” of NRAS-Q61L (The average of Trajectory#1 and Trajectory#2); (D) The comparison of the most dominant conformations of “Average”, “Trajectory#1” and “Trajectory#2”. Red: Switch-I (Average), Orange Red: Switch-I (Trajectory#1), Orange: Switch-I (Trajectory#2), Blue: Switch-II (Average), Cornflower Blue: Switch-II (Trajectory#1), Medium Blue: Switch-II (Trajectory#2), Yellow: P-loop, Green: GDP-bound Mg^2+^. Water and other ions have been hidden.

### Molecular mechanisms for differences in conformational changes between NRAS isoforms

Fig. 14 shows the radial distribution function of "Trajectory#1" and "Trajectory#2". The Fig. 15 shows the averaged free-energy barriers *δW* of two trajectories (T#1 and T#2). For convenience, we list the values of the averaged free-energy barriers *δW* in Table 1. For NRAS-Q61L case, since the hydrophobic pocket and the L61 side chain are non-polar interactions, the time-dependent atomic group–group distances is used here to characterize the interaction between the hydrophobic pocket and the L61 side chain (Fig. 16). At the same time, the The time-dependent atomic site–site distances (Fig. 17) is used to characterize the hydrogen bond interaction between LEU61(H1) and TYP96(O1). This hydrogen bond interaction cooperates with the non-polar interaction of the hydrophobic pocket to firmly fix L61 on the *α*-helix 3, so that the conformational volatility of SW-II is greatly reduced.

**Figure 14:**
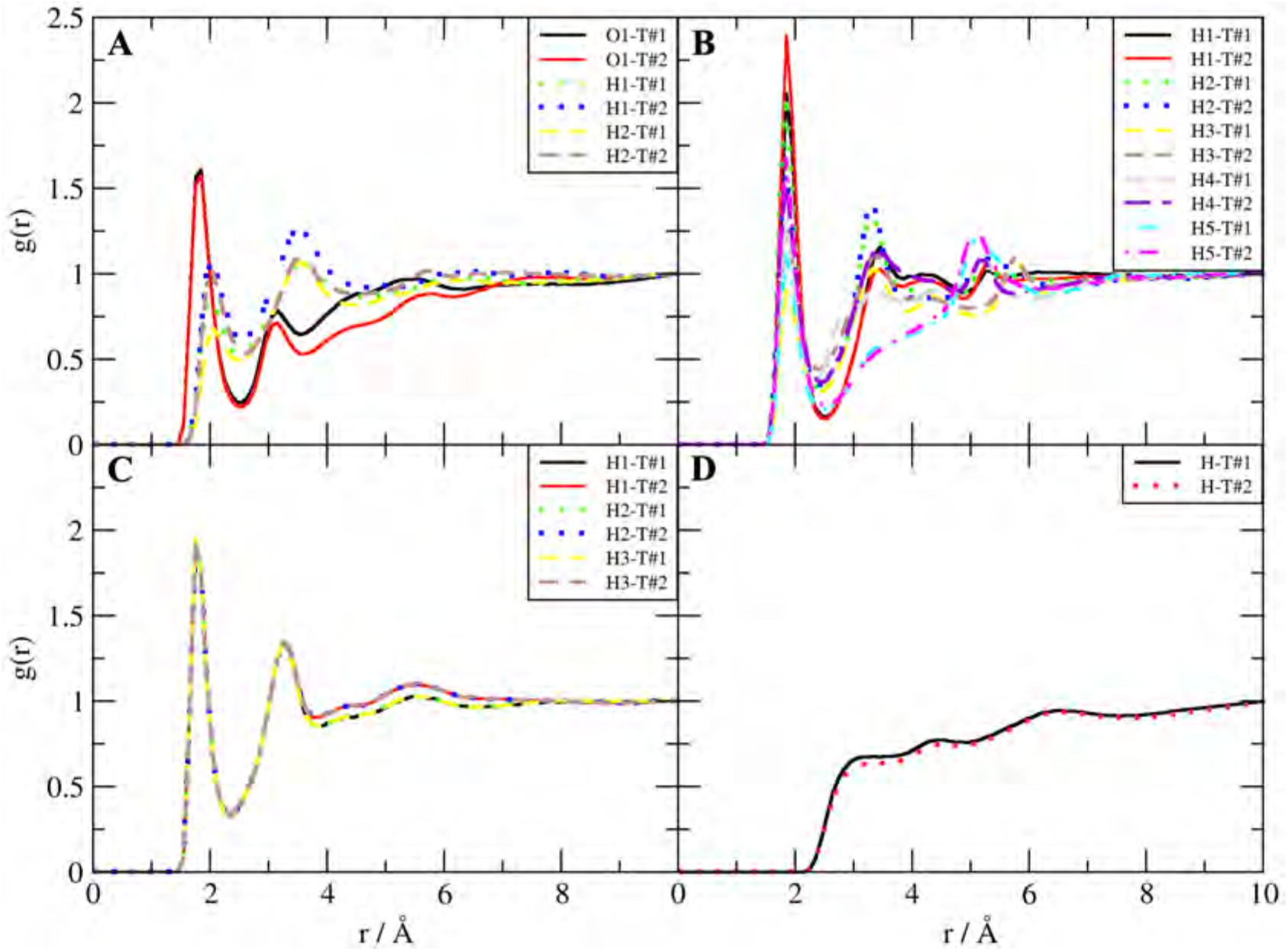
Radial distribution function analysis of the side chain of 61st amino acid of NRAS and water. (A) For NRAS-WT, the radial distribution function analysis of GLN61’s active sites and water; (B) For NRAS-Q61R, the radial distribution function analysis of ARG61’s active sites and water; (C) For NRAS-Q61K, the radial distribution function analysis of LYS61’s active sites and water; (D) For NRAS-Q61L, the radial distribution function analysis of LEU61’s side chain hydrogens and water. T#1 = Trajectory#1, T#2 = Trajectory#2.

**Figure 15:**
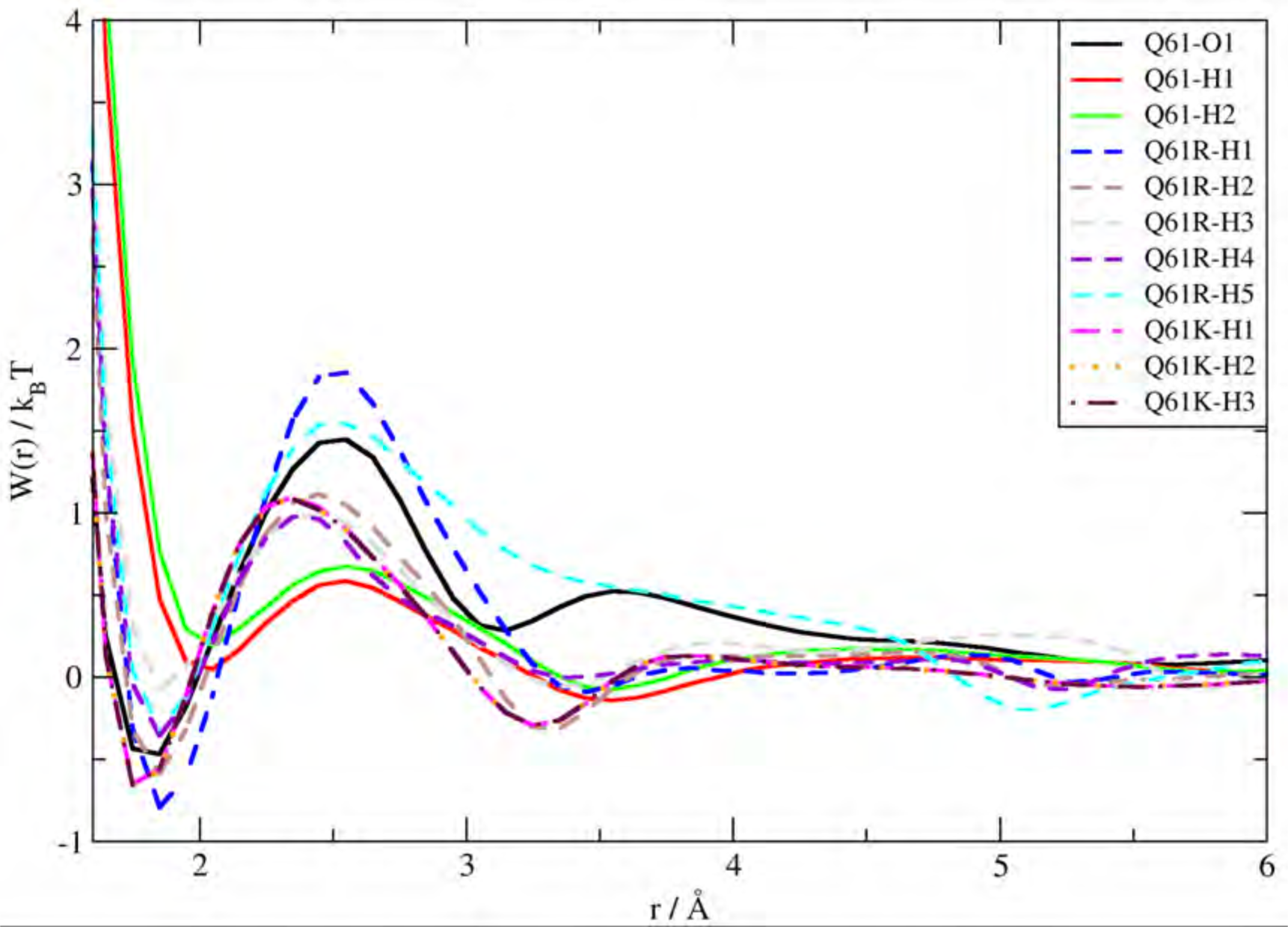
Free-energy barriers *δW* (in *k_B_T*) from reversible work calculations for the interaction between 61st amino acid side chains and water. In order to quantify the height of all barriers, 1 *k_B_T* = 0.616 kcal/mol. *δW* here is the average *δW* after the average of Trajectory#1 and Trajectory#2

**Figure 16:**
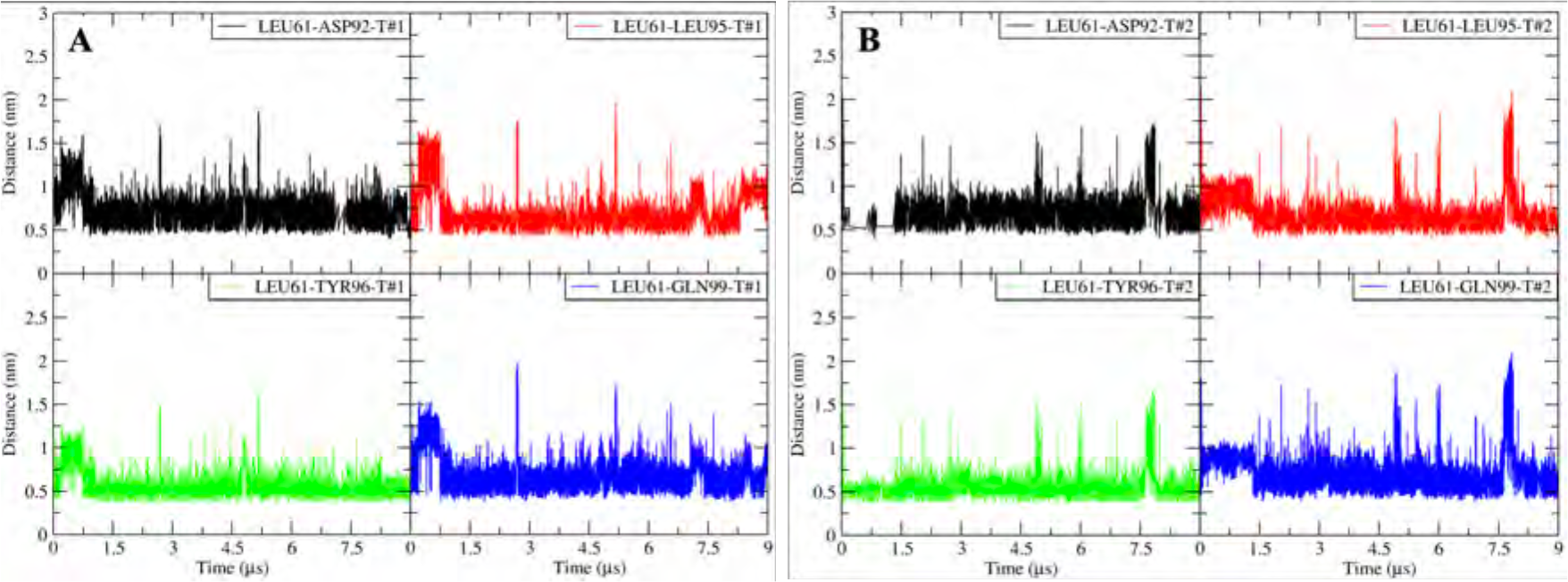
The time-dependent atomic group–group distances between selected amino acid residue side chains, labeled green parts in Fig. 9. (A) Trajectory#1; (B) Trajectory#2. T#1 = Trajectory#1, T#2 = Trajectory#2.

**Figure 17:**
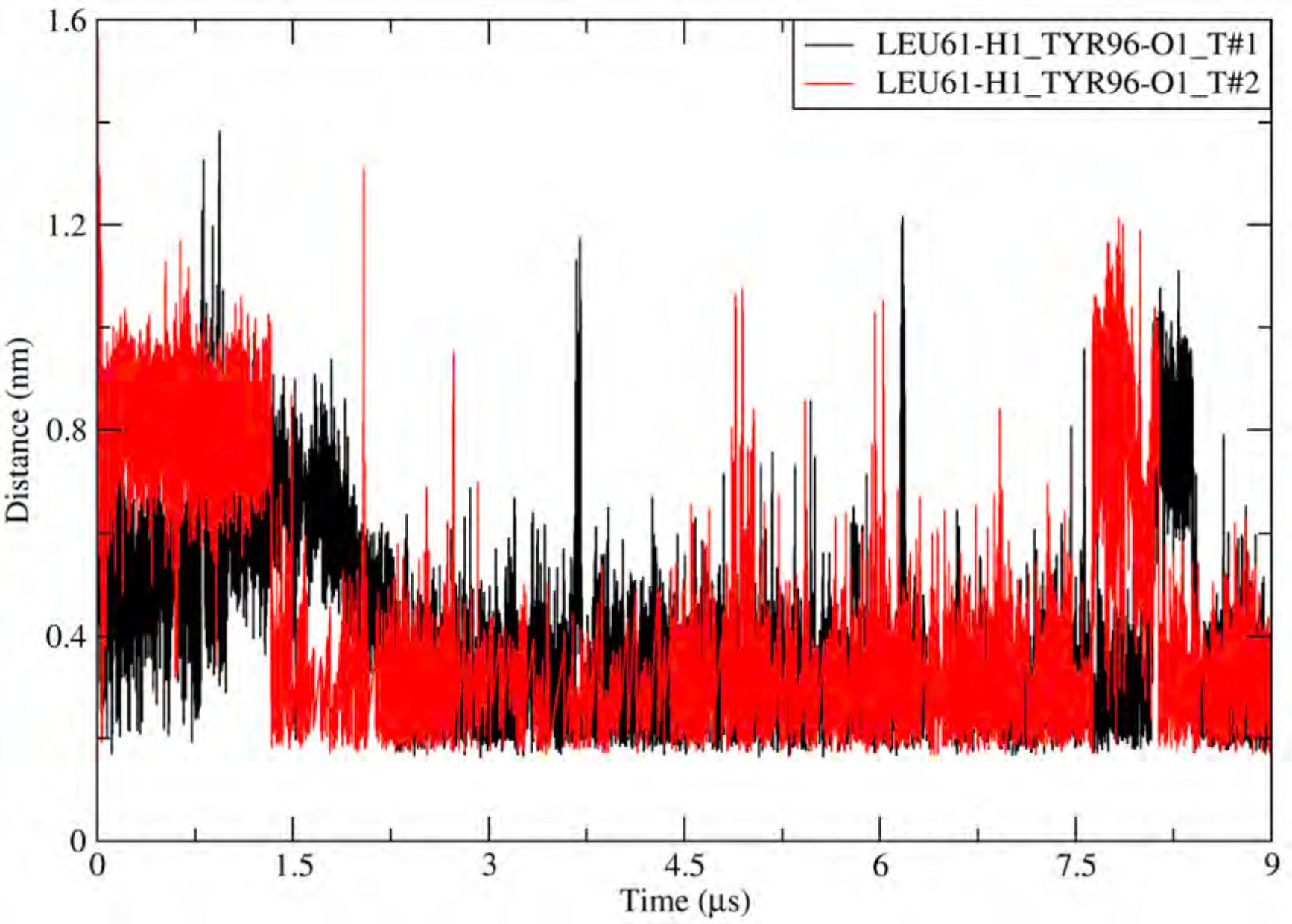
The time-dependent atomic site–site distances between LEU61(H1) and TYP96(O1).

### Details in the drug design process

Here we take **MRTX1133** as the isomer-sourced template in our **ISSI** drug design process. And the receptor structure comes from the NRAS MD simulation results in the first part of this paper. The Fig. 18 shows the targetable pocket of NRAS-Q61R, to better visualize the shape of the pocket and the distribution of different atoms (such as white H atoms, red O atoms and blue N atoms) on its surface, we used the "heteroatom" coloring method of "Chimera" software. The targetable pocket of NRAS-Q61R is mainly composed of three parts, as shown in Fig. 18A: green box **Ia**, cyan circle **IIa** and orange box **IIIa**. As we can see in green box **Ia** surface area, there are many polar atoms distributed on the surface of box **Ia**, such as hydrogen bond acceptor oxygen atom and nitrogen atom. In contrast to green box **Ia** area, cyan circle **IIa** area is a hydrophobic pocket located deep within the NRAS protein (near the P-loop). As for orange box **IIIa** area, this part is located between *α*-helix 2 and *α*-helix 3, has a more open shape and a balanced distribution of polar atoms compared to green box **Ia** area.

**Figure 18:**
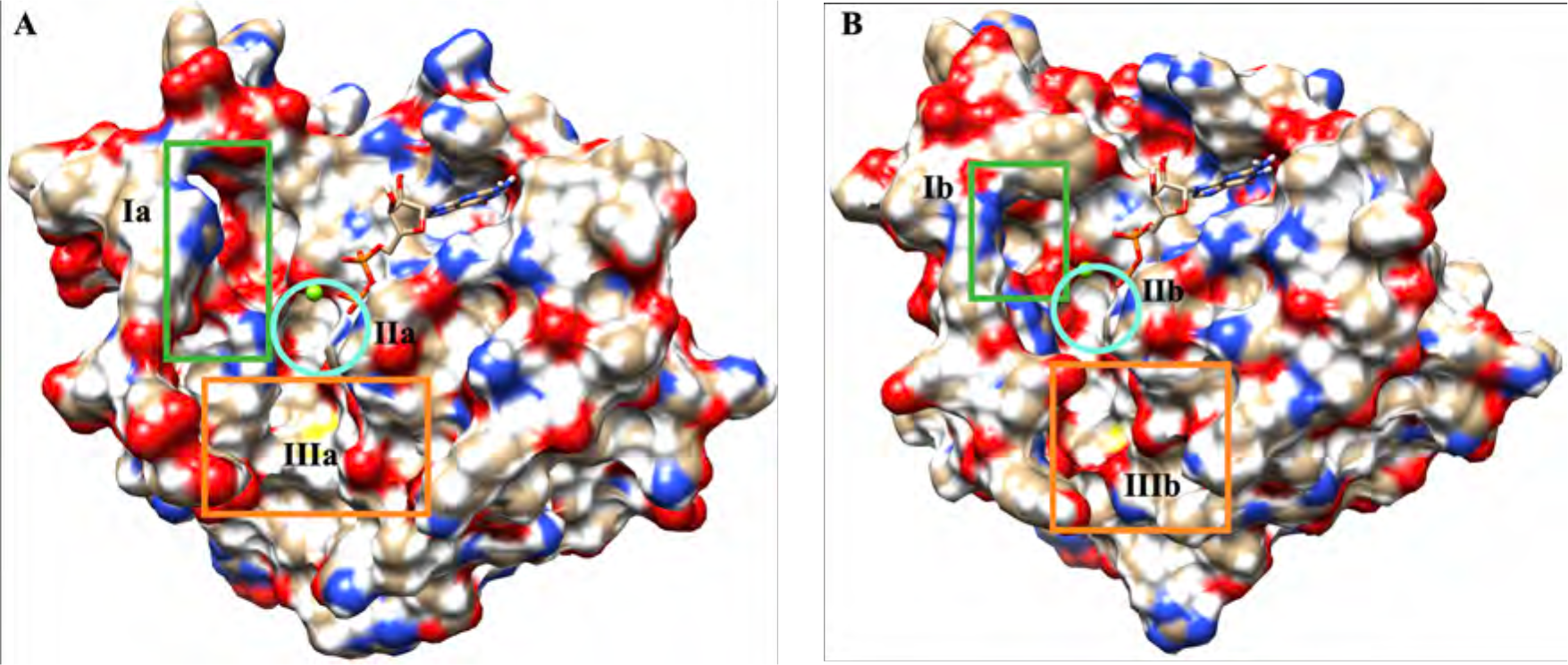
Representative conformations (NRAS-Q61R) used in the **ISSI** drug design approach. (A) The SW-II has a larger opening; (B) The SW-II holds a smaller opening.

**Figure 19:**
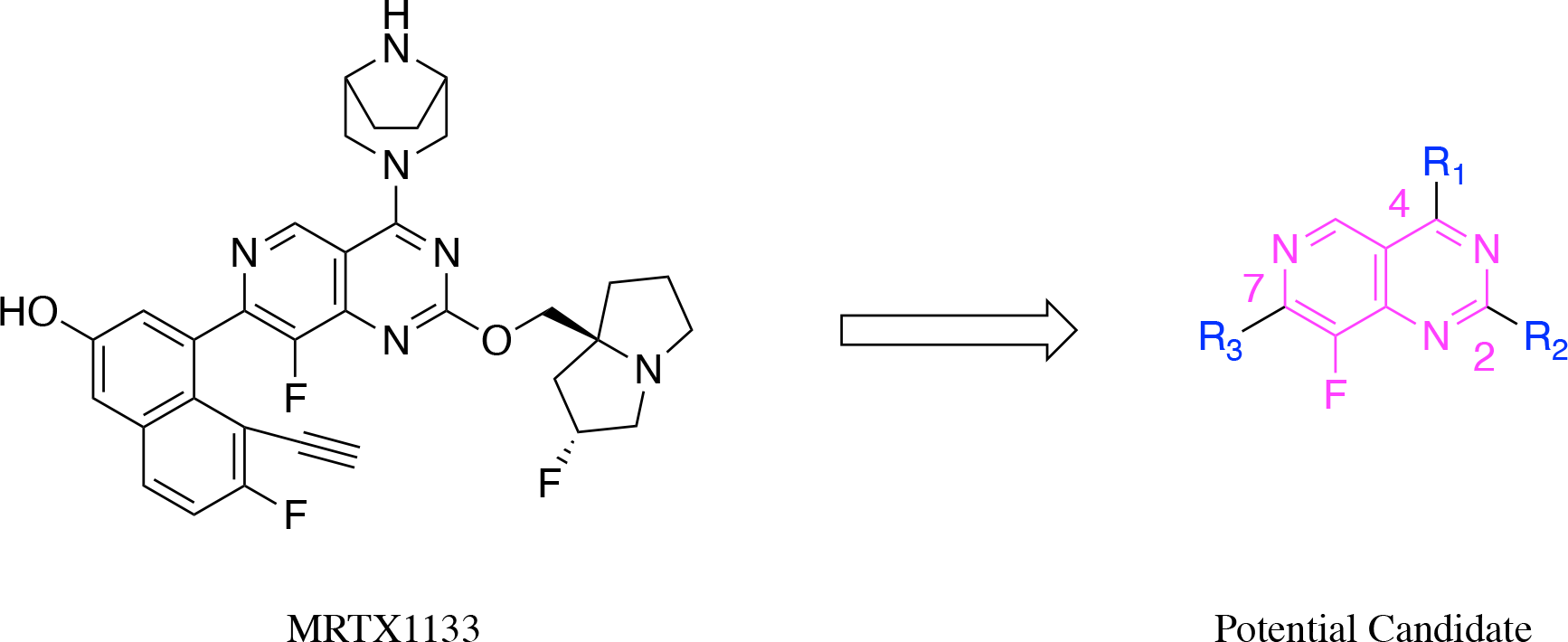
The structure of MRTX1133 and the pyrido[4,3-*d*]pyrimidine core for further optimization.

After determining the "isomer-sourced template" and "receptor structure", we mainly carried out three generation of structural iteration, and the corresponding structural iteration data and screening process can be seen from Fig. 20 to Fig. 24. Firstly, the "AutoDock" analysis tool^49^ was employed for measuring the compatibility of MRTX1133 with the pocket located on the surface of NRAS-Q61R, see Fig. 20 and Fig. 21 B. The compound **A1** (MRTX1133) with a [3.2.1]bicyclic diamino substituent had the docking score from -7.5 to -9.1 and formed 2 HB with the NRAS-Q61R pocket. More importantly, the [3.2.1]bicyclic diamino group at the C4-position prefers to bind to the green box **Ia** area of the NRAS-Q61R pocket. As we know, there are many hydrogen-bonding acceptor atoms distributed in the green box **Ia** area, which is very favorable for forming hydrogen-bonding interactions with potential inhibitor molecules. And the [3.2.1]bicyclic diamino group located on C4-position was originally designed to form a salt bridge with ASP12 of KRAS-G12D.

**Figure 20:**
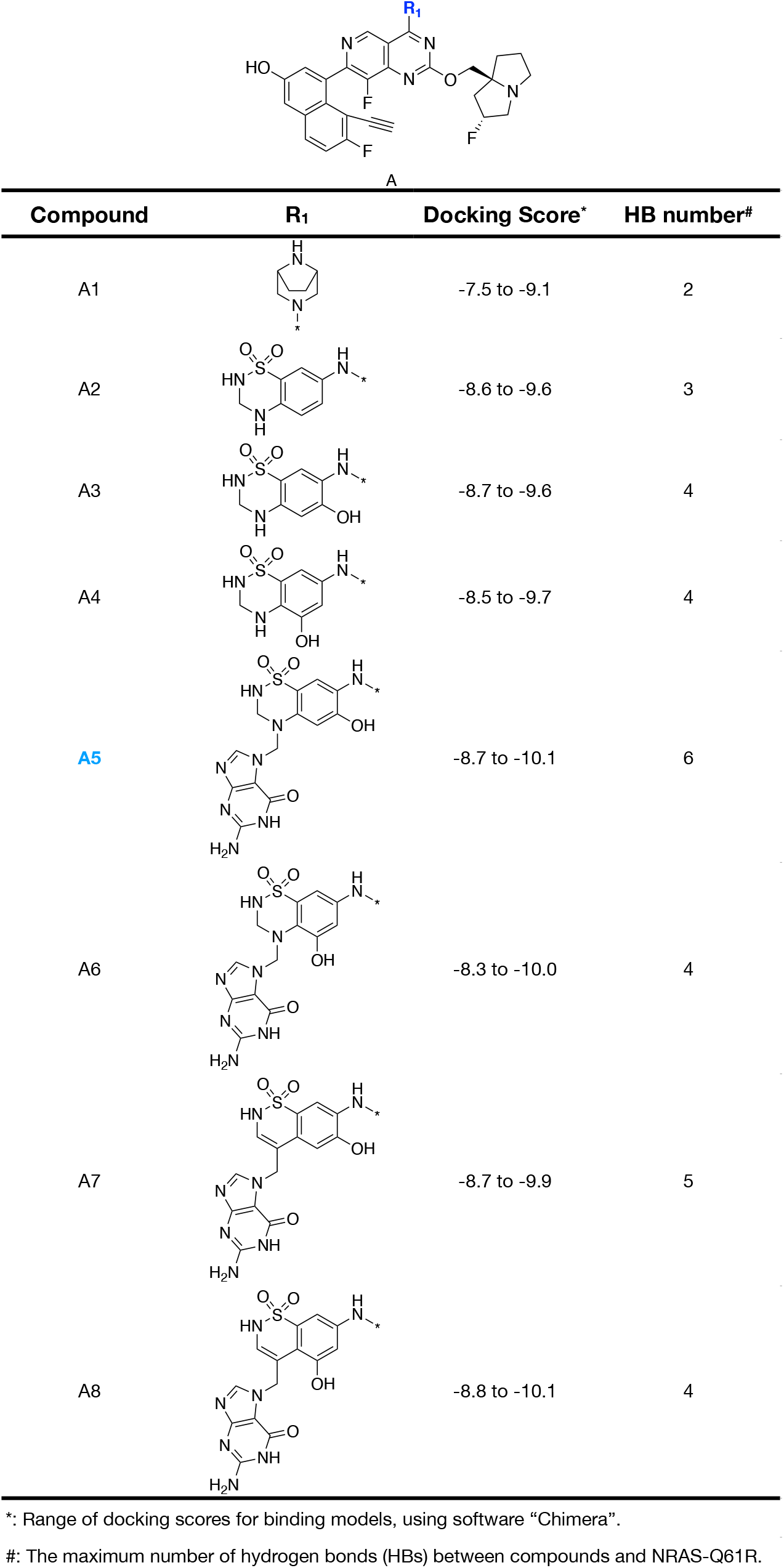
The structural iteration and optimization of substituents (*R*_1_) at the C4-position of pyrido[4,3-*d*]pyrimidine core. Based on the Fig. 18-A.

**Figure 21:**
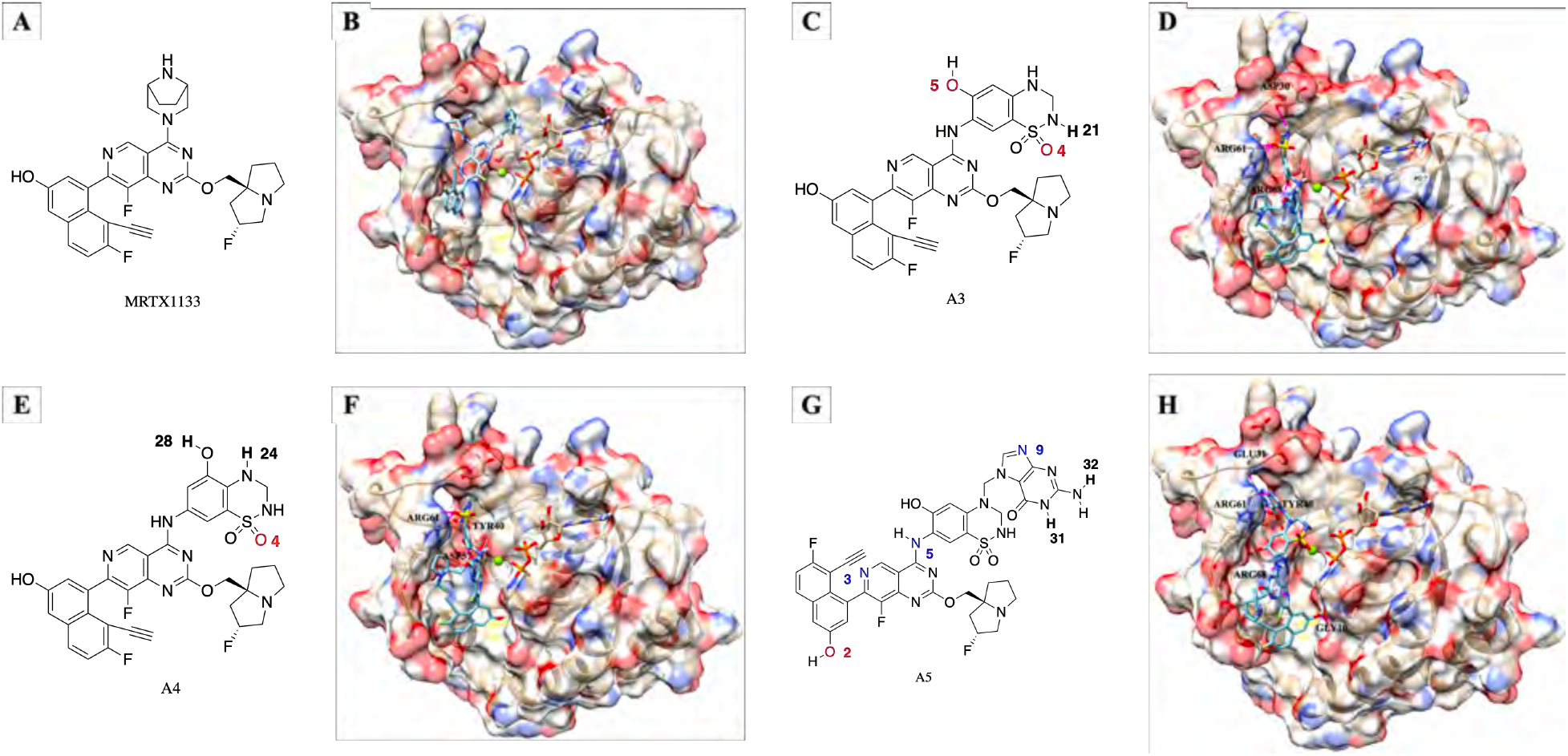
Screenshots of representative compounds and their protein binding modes during structure iteration and optimization of substituent *R*_1_. Based on the Fig. 18-A.

**Figure 22:**
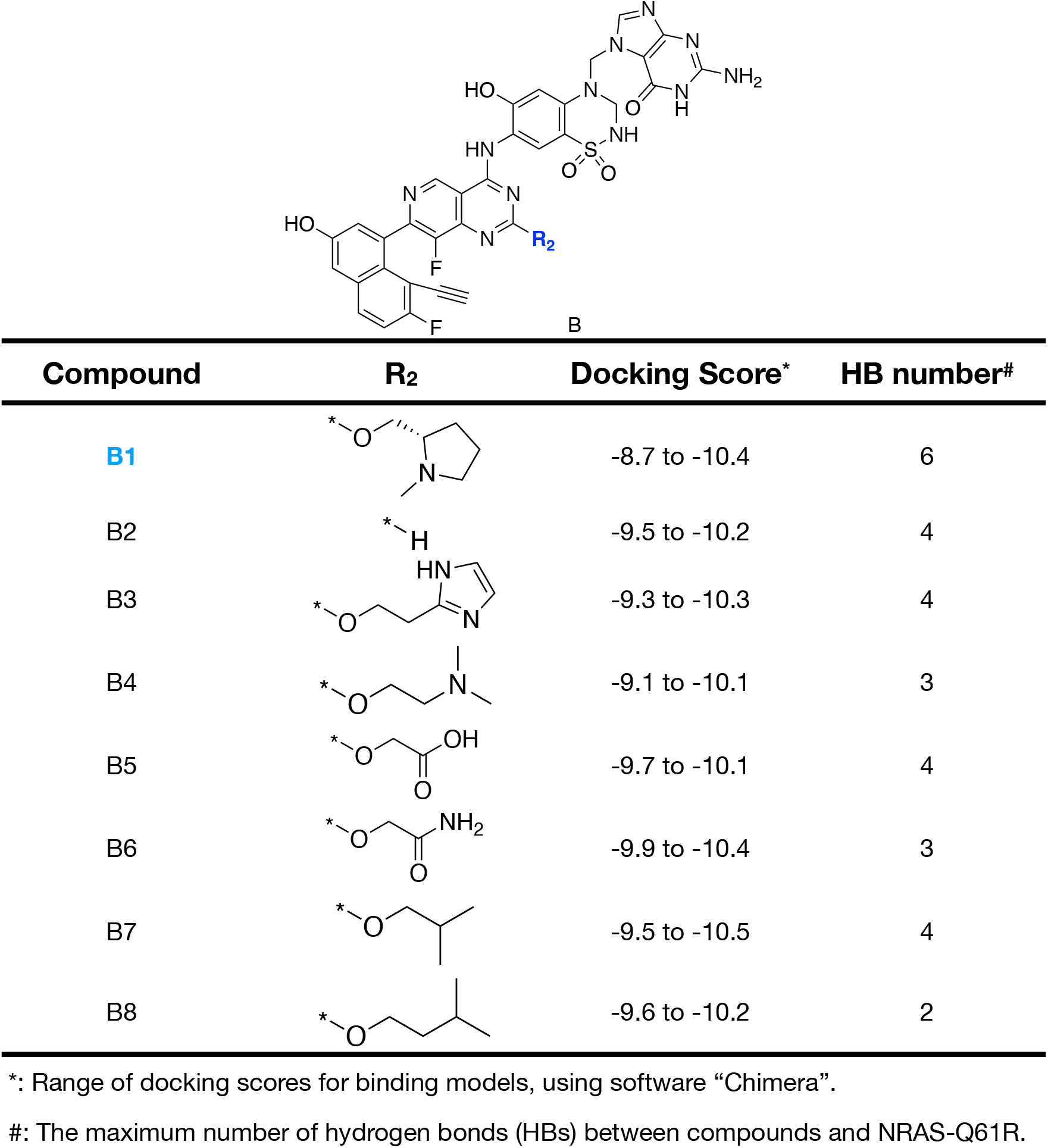
The structural iteration and optimization of substituents (*R*_2_) at the C2-position of pyrido[4,3-*d*]pyrimidine core. Based on the Fig. 18-A.

**Figure 23:**
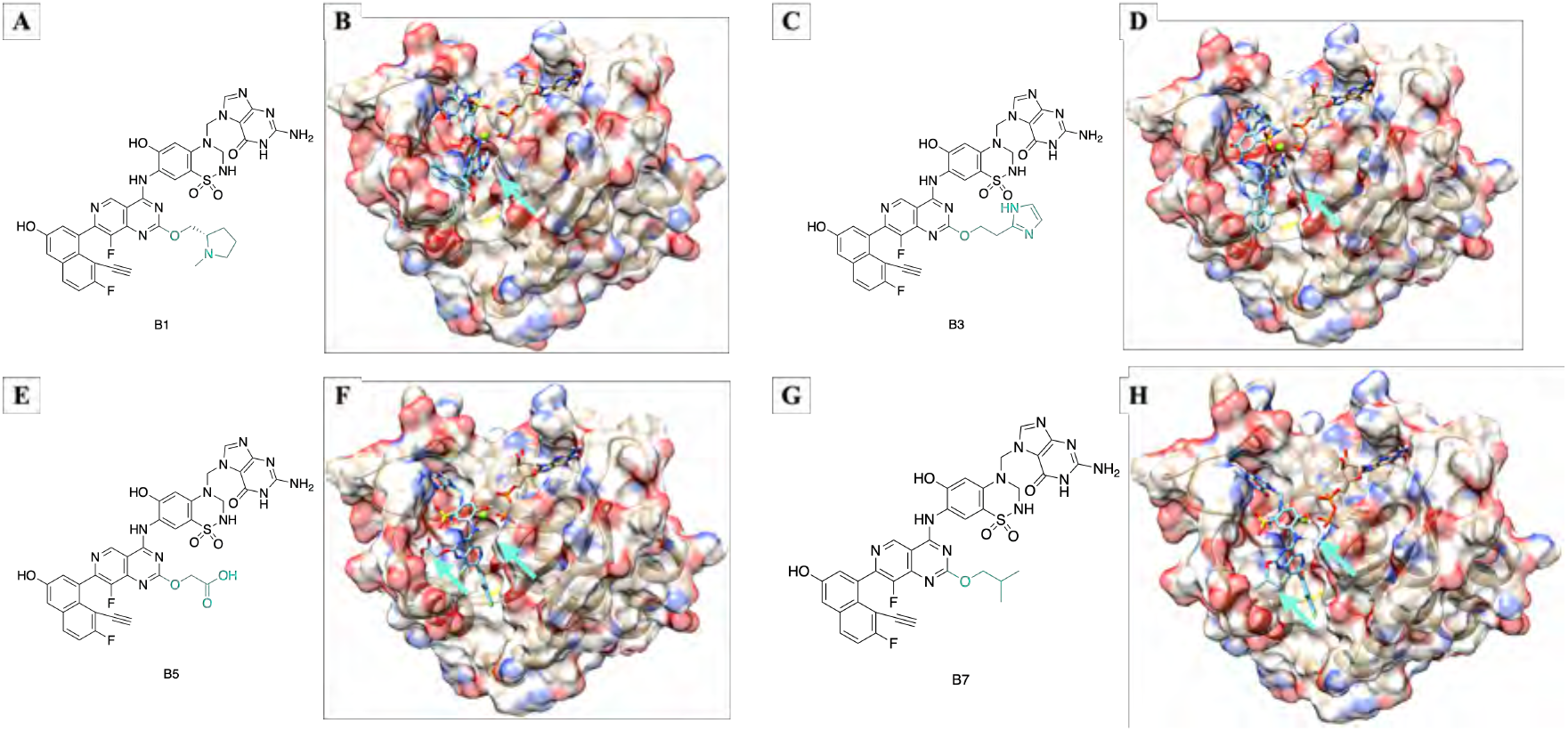
Screenshots of representative compounds and their protein binding modes during structure iteration and optimization of substituent *R*_2_. Based on the Fig. 18-A.

**Figure 24:**
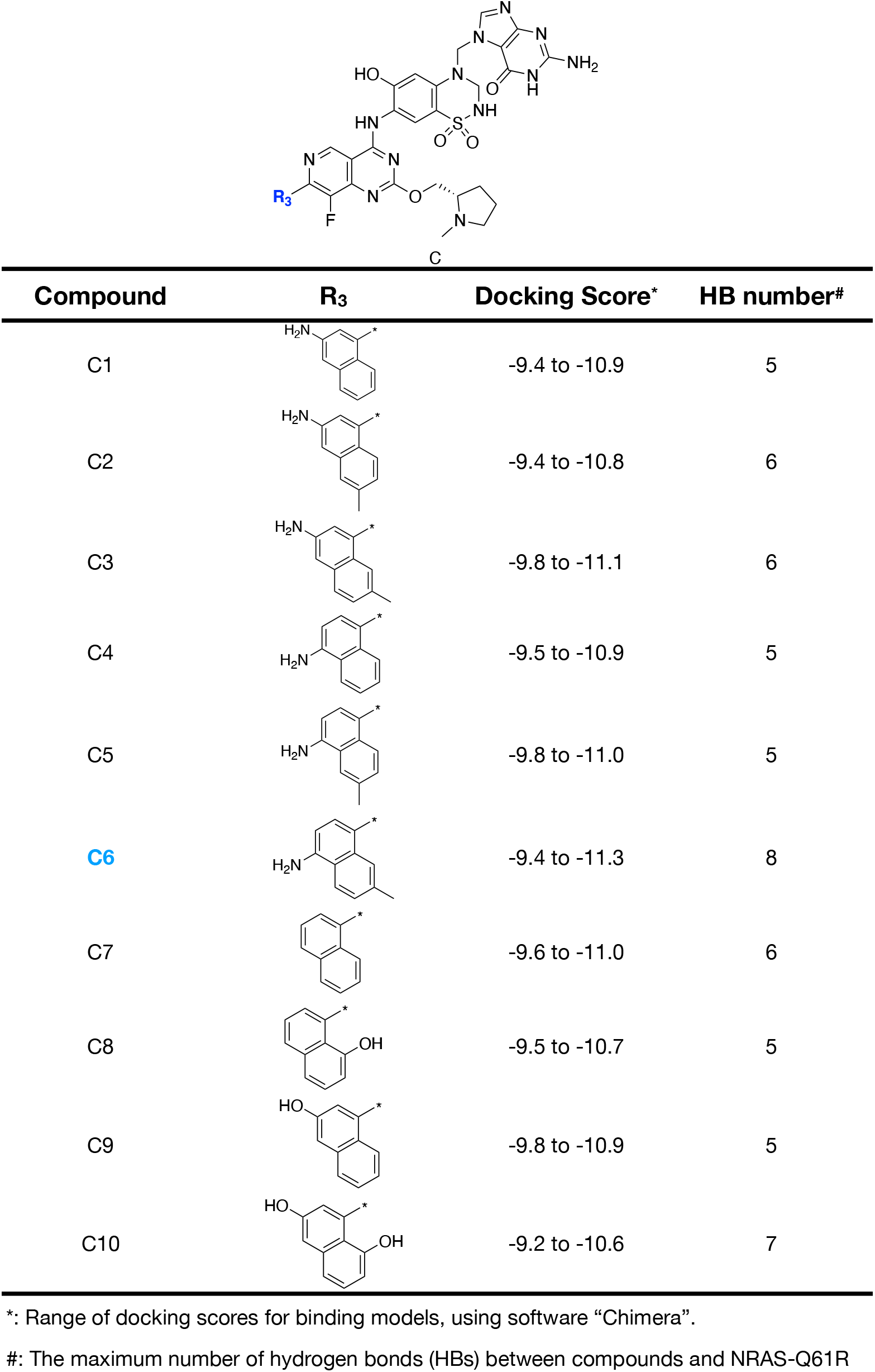
The structural iteration and optimization of substituents (*R*_3_) at the C7-position of pyrido[4,3-*d*]pyrimidine core. Based on the Fig. 18-A.

Given this valuable information, we began optimization of the series at the C4-position keeping the C2-pyrrolizidine and C7-naphthol substituent unchanged (Fig. 20). The docking score of compound **A2** with the DBD group was around -8.6 to -9.6 and the HB interaction with NRAS-Q61R is slightly enhanced compared with compound **A1**. When the phenolic hydroxyl group was added in the benzene ring of DBD substituent, the HB interaction between compounds **A3-A4** and NRAS-Q61R continued to strengthen, which was 2-fold larger than compound **A1**. For instance, the H21 atom of **A3** form HB with backbone carbonyl oxygen atom of ASP30, the sulfonyl oxygen O4 of compound **A3** form 2 HBs with ARG61 side chain amino group and the oxygen atom O5 of the phenolic hydroxyl group form HB interaction with the side chain of ARG68, Fig. 21 C-D. While change the position of phenolic hydroxyl group in the benzene ring (Fig. 21 E), the compound **A4** exhibited similar properties to **A3**, the H24 of **A4** form HB with side chain of TYR40, the sulfonyl oxygen O4 of compound **A4** form 2 HBs with ARG61 side chain amino group and the phenolic hydroxyl group (H28) form HB interaction with side chain carboxyl group of ASP57, Fig. 21 F. These observations further confirmed that introducing a substituent with the active site for HB interaction at C4-position of pyrido[4,3-*d*]pyrimidine core is a correct direction. Further optimization (**A5-A8**) revealed the introduction of guanine module into the DBD substituent at the C4-position is of great benefit in improving the affinity for the targetable pocket of NRAS-Q61R. Especially for compound **A5**, its docking score reached the maximum value of this optimization, and the strength of the HB interaction with the surface pocket of NRAS-Q61R was 3-fold larger than compound **A1** and 1.5-fold larger than that of **A3-A4**, Fig. 20. According to the the docking results of **A5** and NRAS-Q61R (Fig. 21 G-H), firstly, the guanine module matches the spatial structure of green box **Ia** and can establish stable interactions with it, and secondly, the existence of the guanine group expands the length of the molecule, making the other active atoms on **A5** can establish more interaction with the druggable pocket of the NRAS-Q61R. For instance, the atoms on the guanine group (N9, H31 and H32) can form HB interactions with the ARG61 side chain amino hydrogen atom, TYR40 phenolic hydroxyl oxygen atom and GLU31 backbone carbonyl oxygen atom, respectively. At the same time, the N3 and N5 atoms of compound **A5** form HB interactions with ARG68 side chain amino group, and the phenolic hydroxyl oxygen (O2) of compound **A5** also forms HB interactions with the GLY10 backbone amide bond hydrogen atom (Fig. 21 H).

With a promising C4-substitution in hand, we next focused on optimizing the C2-position of the pyrido[4,3-*d*]pyrimidine core, which is summarized in Fig. 22. Inspired by the docking results of the C4-position optimization process, we noticed that the original C2-pyrrolizidine always tended to move away from the cyan circle **IIa** direction during the C4 optimization, Fig. 21 D, F and H. Considering pyrrolizidine is bicyclic compound with a large volume together with the circular hydrophobic pocket cyan circle **IIa** region, so we first tested the compound **B1** substituted by pyrrolidinyl. The docking results of compound **B1** with NRAS-Q61R shows that compared with compound **A5**, the affinity of **B1** to NRAS-Q61R was further improved and C2-pyrrolidinyl was fit well for the cyan circle **IIa** region, Fig. 23B. Deletion of the C2-pyrrolizidine substitution in **A5** yielded compound **B2**, which resulted in the decrease of HB forming potency, confirming the importance of this moiety for the overall affinity of the designed compound. The imidazole group has a similar size to the pyrrolidinyl group (Fig. 23D), but the HB forming potency of compound **B3** is weaker than that of **B1**. In addition to the cyclic substituents mentioned above, the chain substituents at the C2-position were also under consideration. Such as *N,N* -dimethylethanamine group (compound **B4**), carboxyl (compound **B5**), acetamide substituent (compound **B6**), isobutane substituent (compound **B7**) and isopentane substituent (compound **B8**). But none of them were more potent than the pyrrolidinyl analog **B1**. In contrast, some compounds with chain substituents have the dominant docking conformation where the C2-substituent is located away from the cyan circle **IIa** region region, Fig. 23F and H. This C2-position optimization revealed the pyrrolidinyl was benefit for improving the binding affinity.

Further optimization was displayed on the C7-position of pyrido[4,3-*d*]pyrimidine core using the naphthyl amine and naphthol substituents, Fig. 24. For this exploration, the docking score of compound **C7** with NRAS-Q61R was further improved when using the unsubstituted naphthyl, which means that the structural fit of compound **C7** to the targetable pocket of NRAS-Q61R was further enhanced. Simple naphthyl monosubstitutions such as 3-amino, 4-amino, 8-phenolic hydroxyl or 3-phenolic hydroxyl (Fig. 24, compound **C1**, **C4**, **C8** and **C9**) provided similar docking scores and slightly lower HB forming ability than unsubstituted **C7**. Next is the screening of naphthyl disubstituents, as for 6-methylnaphthalen-3-amine (compound **C2**), 7-methylnaphthalen-3-amine (compound **C3**) and 6-methylnaphthalen-4-amine (compound **C5**) also hold the similar activity to **C7**. The Compound **C10** with naphthalene-3,8-diol as the substituent has further improved affinity to the targetable pocket of NRAS-Q61R. Finally, the compound **C6** with 7-methylnaphthalen-4-amine as the substituent hit the goal, which holds both the highest docking score and the strongest HB forming ability. Considering that the conformation of the protein receptor has an important impact on the docking analysis,^50,51^ another conformation of NRAS-Q61R (Fig. 18 B) was used as the receptor for another round of ISSI drug design, see Fig. 25 to Fig. 27. The size of the targetable pocket is reduced in conformation B compared to conformation A, especially in region **Ib**. At the same time, the docking results showed that the HB formation ability of the screened compounds was slightly weakened, but the results of the two rounds of screening finally pointed to compound **C6** (**HM-516**).

**Figure 25:**
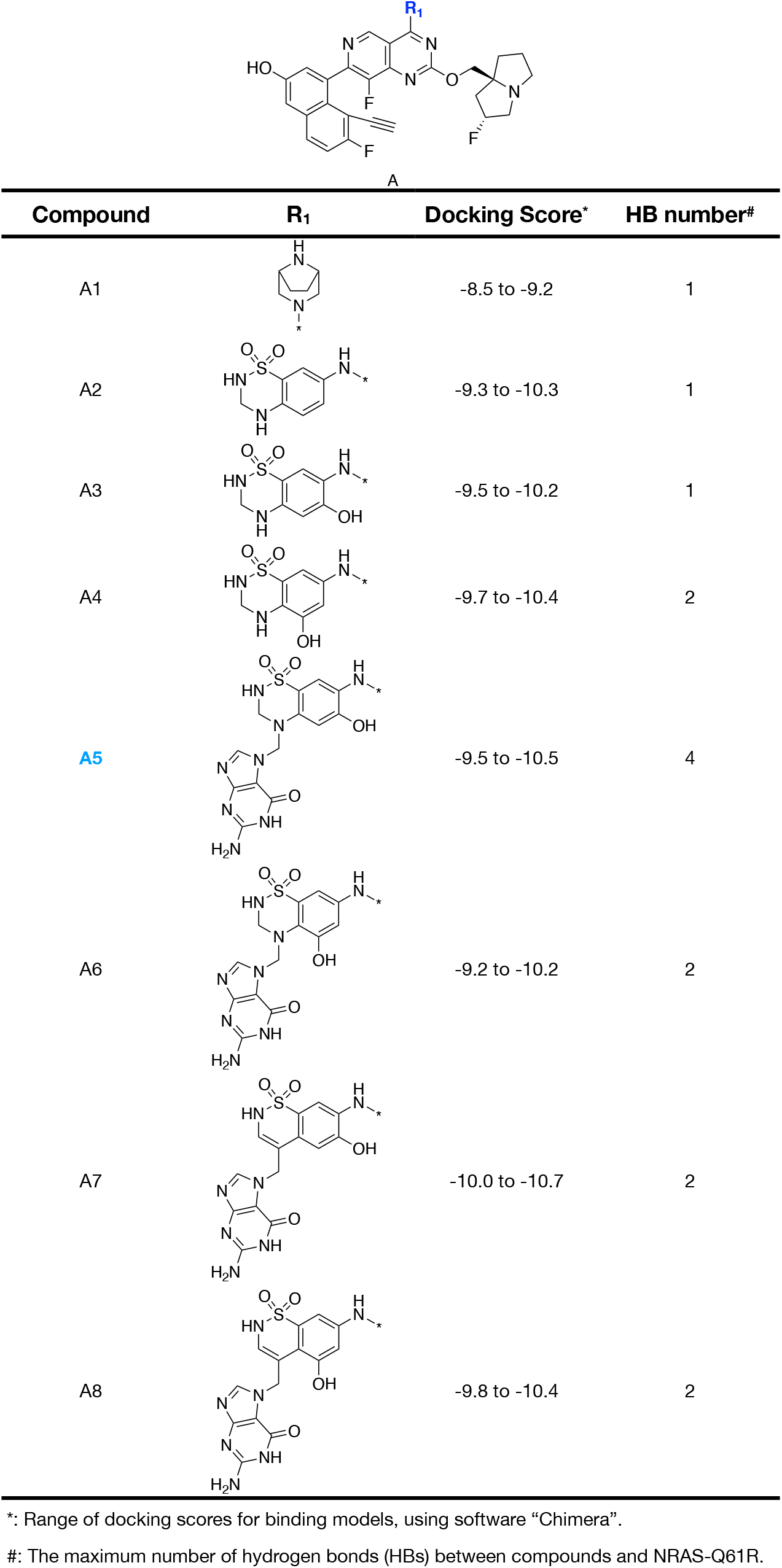
The structural iteration and optimization of substituents (*R*_1_) at the C4-position of pyrido[4,3-*d*]pyrimidine core. Based on the Fig. 18-B.

**Figure 26:**
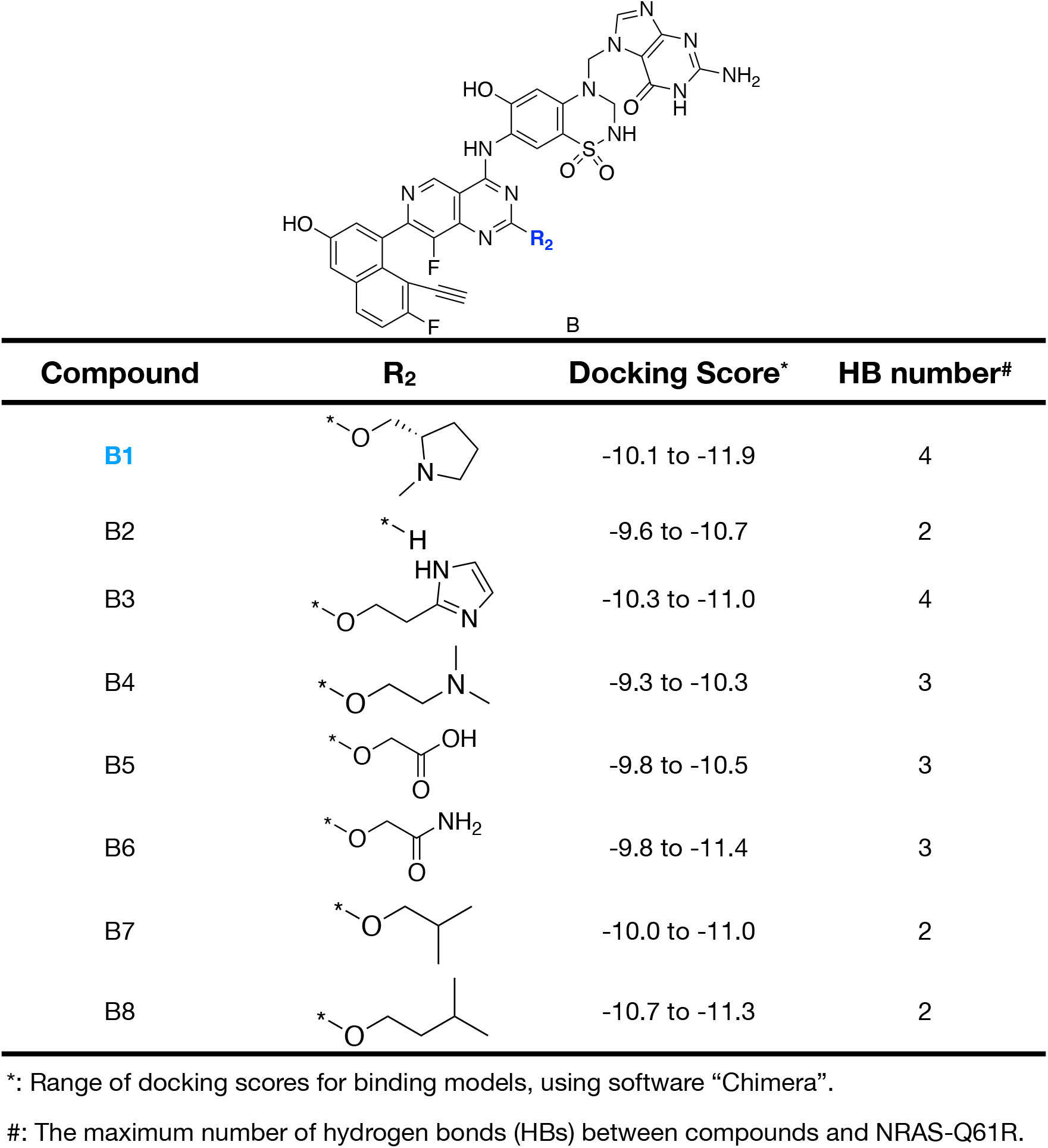
The structural iteration and optimization of substituents (*R*_2_) at the C2-position of pyrido[4,3-*d*]pyrimidine core. Based on the Fig. 18-B.

**Figure 27:**
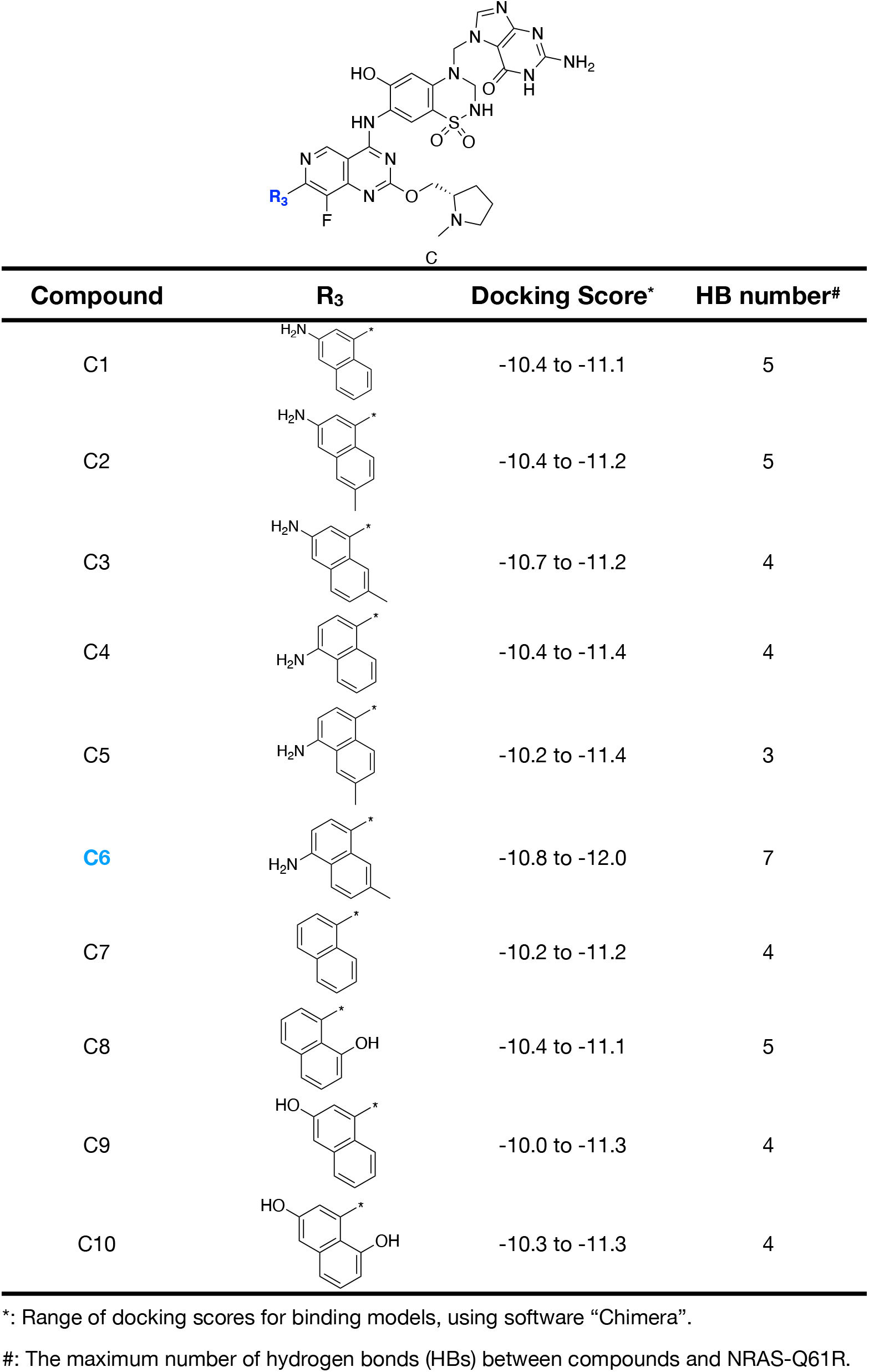
The structural iteration and optimization of substituents (*R*_3_) at the C7-position of pyrido[4,3-*d*]pyrimidine core. Based on the Fig. 18-B.

Afterwards, four long independent trajectories (total 10 *µ*s) have been produced for exploring the dynamic binding mode of **HM-516** to NRAS-Q61R, Fig. 28. The "Gibbs Free-Energy Landscape" of the individual trajectories in Fig. 28 nicely demonstrates the convergence of our results. During total of 10 *µ*s of simulation, the process of drug association and dissociation was monitored. In Fig. 28, the energy wells in the blue circle represent the state where the **HM-516** is tightly bound to NRAS-Q61R surface druggable pocket. The energy well in the red circle represents the dissociation state of the drug from NRAS-Q61R, while the energy wells in the purple circles represent a state between tight association and dissociation. Theoretically, when the situation time tends to infinity, the four "Gibbs Free-Energy Landscape" in Fig. 28 will tend to be consistent. Moreover, we also provide corresponding snapshots of different binding states of the **HM-516** to NRAS-Q61R, see Fig. 29.

**Figure 28:**
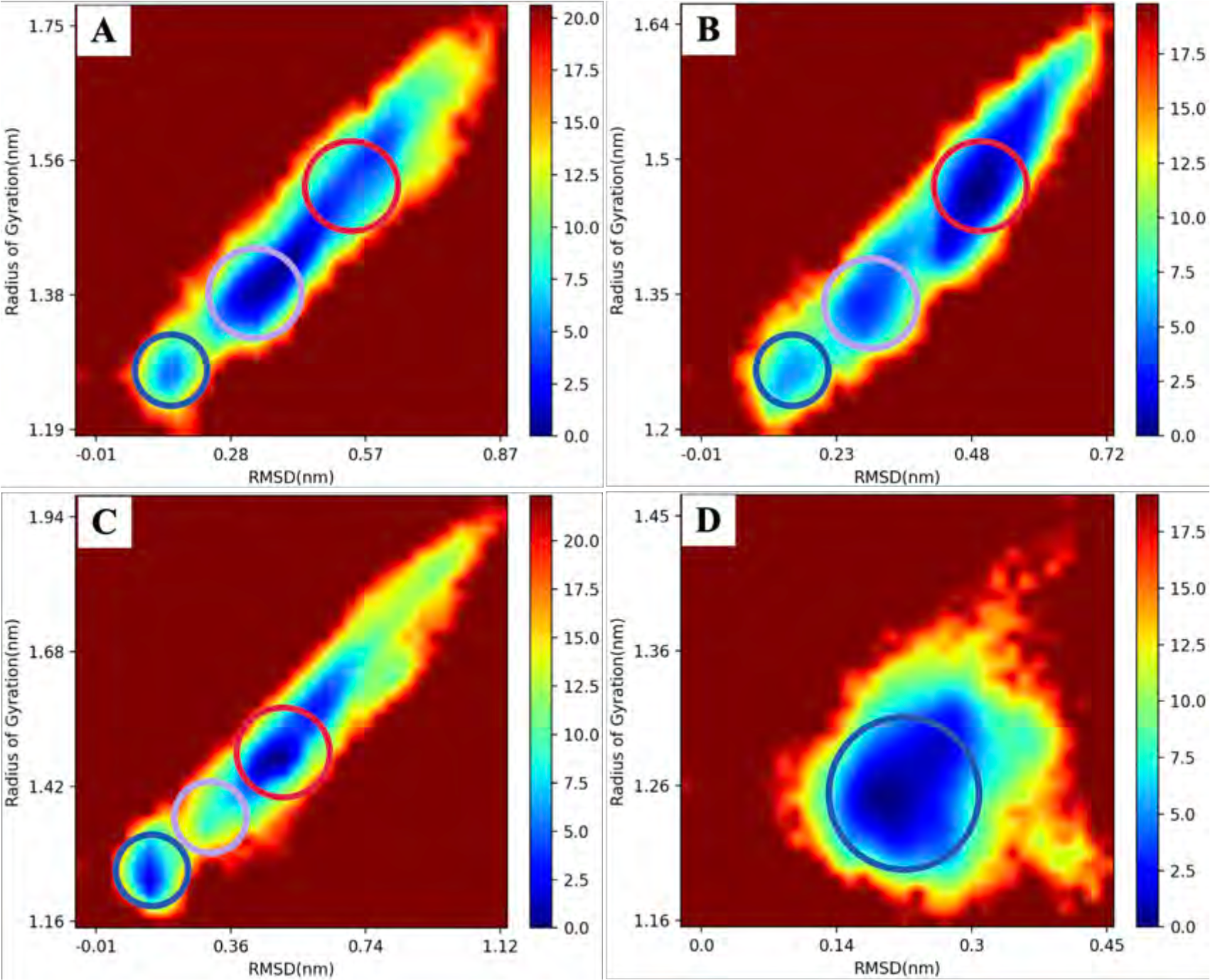
The "Gibbs Free Energy" analysis of NRAS-Q61R simulation together with 10 **HM-516**. (A) Trajectory#1, total 2 *µ*s; (B) Trajectory#2, total 2 *µ*s; (C) Trajectory#3, total 2 *µ*s; (D) Trajectory#4, total 4 *µ*s.

**Figure 29:**
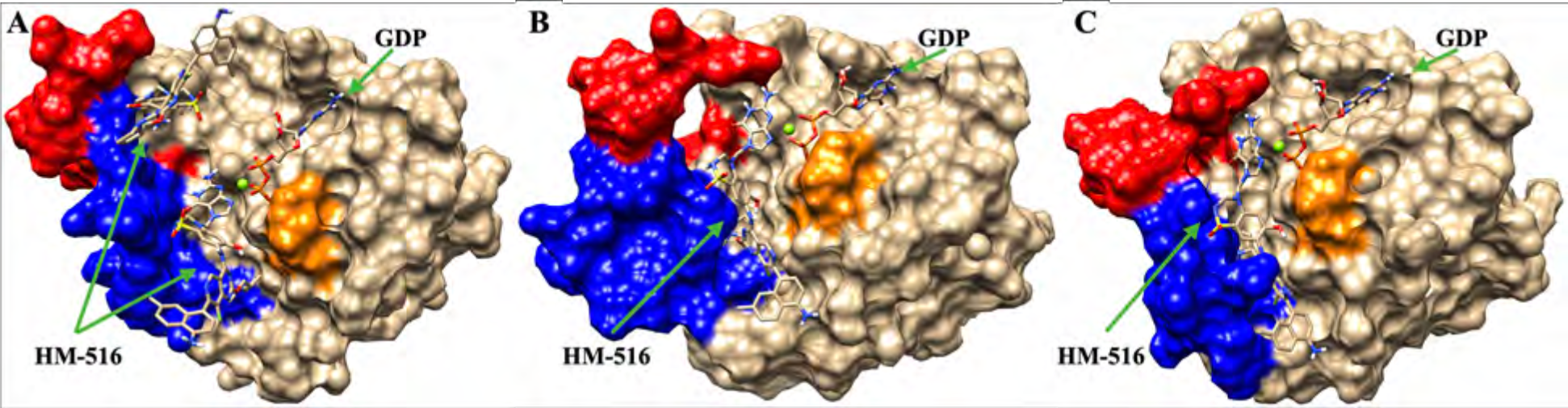
The corresponding snapshots of different binding states of the **HM-516** to NRAS-Q61R. (A)The dissociation state of the **HM-516** from NRAS-Q61R; (B) The intermediate binding state of **HM-516** and NRAS-Q61R, between dissociation and tight association; (C) The tight binding status of **HM-516** to NRAS-Q61R. The rest of the **HM-516** are hidden.

## Acknowledgement

Please use “The authors thank . . . ” rather than “The authors would like to thank . . . ”.

The author thanks Mats Dahlgren for version one of achemso, and Donald Arseneau for the code taken from cite to move citations after punctuation. Many users have provided feedback on the class, which is reflected in all of the different demonstrations shown in this document.

## Supporting Information Available

A listing of the contents of each file supplied as Supporting Information should be included. For instructions on what should be included in the Supporting Information as well as how to prepare this material for publications, refer to the journal’s Instructions for Authors.

The following files are available free of charge.

- Filename: brief description
- Filename: brief description

